# Adaptive thermogenesis is mediated by GDF15 via the GFRAL neuronal axis in mice

**DOI:** 10.1101/2024.01.15.575796

**Authors:** Ji Eun Kim, Sang-Hyeon Ju, Min Hee Lee, Hyun Jung Hong, Uzma Yaseen, Jung Tae Kim, Benyuan Zhang, Hyon-Seung Yi, Seong Eun Lee, Yea Eun Kang, Yoon-Sun Yi, Sangmi Jun, Minsung Park, Jinkuk Kim, Johan Auwerx, Jong-Woo Sohn, Ju Hee Lee, Minho Shong

**Affiliations:** Research Center for Endocrine and Metabolic Diseases, School of Medicine, Chungnam National University, Daejeon 35015, Republic of Korea; Graduate School of Medical Science and Education, Korea Advanced Institute of Science and Technology, 291 Daehak-ro, Yuseong-gu, Daejeon 34141, Republic of Korea; THOR Therapeutics, Inc., 99 Chungnam National University, Daehak-ro, Yuseong-gu, Daejeon 34134, Republic of Korea; Department of Internal Medicine, Chungnam National University Hospital, Daejeon, 35015, Republic of Korea; Center for Research Equipment, Korea Basic Science Institute, Cheongju 28119, Republic of Korea; Laboratory for Integrative Systems Physiology, Institute of Bioengineering, École Polytechnique Fédérale de Lausanne, Lausanne 1015, Switzerland

**Keywords:** Brown adipose tissue, Glial cell-derived neurotrophic factor receptor alpha-like neuron, Growth differentiation factor 15, Mitochondrial unfolded protein response, sympathetic nervous system

## Abstract

Adaptive thermogenesis is a key homeostatic mechanism that primarily occurs in brown adipocytes and enables the maintenance of body temperature. Although this process involves coordinated responses in multiple tissues, including the browning of white adipocytes, the precise inter-organ crosstalk underlying adaptive thermogenesis is unclear. Here, we investigate the pivotal role of the GDNF family receptor alpha-like (GFRAL) neuronal axis in modulating compensatory thermogenic responses in brown and white adipose depots under stress conditions, specifically the mitochondrial unfolded protein response resulting from genetic modification and cold exposure. We employed a mouse model with targeted deletion of *Crif1* in the mitoribosomes of brown adipocytes, and cold-exposed mice and immortalized adipocytes, to uncover the mechanism by which mitochondrial stress-induced growth differentiation factor 15 (GDF15) expression affects metabolism and facilitates adaptive thermogenesis. We found that *Crif1* deletion resulted in browning of inguinal white adipose depots, increased energy expenditure, reduced food intake, and resistance to weight gain. Retrograde neuronal tracing established that GFRAL-positive neurons in the hindbrain and sympathetic preganglionic neurons in the spinal cord mediated the GDF15-associated browning of inguinal white adipose tissue. Intervention studies using antisense oligonucleotides to inhibit *Gfral* expression blunted the effect of *Crif1* deletion on energy expenditure and food intake, further confirming the essential role the GFRAL axis plays in GDF15-driven thermogenic adaptation in white adipose tissue. Our findings suggest that the GFRAL neuronal axis is key in coordinating the adaptive thermogenic response across multiple tissues and adipose depots, thereby ensuring metabolic homeostasis during mitochondrial stress.

## Introduction

Adaptive thermogenesis in adipose tissue is a crucial homeostatic mechanism that helps protect animals against cold environments. While brown adipose tissue (BAT) is the primary site of thermogenesis, beige adipocytes within white adipose tissue (WAT) also display a thermogenic response to cold-induced hypothermia. The coordinated actions of brown and beige adipocytes, facilitated by activation of the sympathetic nervous system (SNS), are essential for the regulation of body temperature in mammals (Cannon & Nedergaard, 2004). The SNS not only triggers lipolysis in white adipocytes, supplying substrates for brown adipocyte thermogenesis, but also appears to orchestrate the temporary emergence of beige adipocytes, indicating a complex regulatory hierarchy underlying responses to cold in adipose tissue (Nedergaard & Cannon, 2014).

In circumstances where BAT is absent or functionally compromised, WAT has the capacity to undergo compensatory browning, a process driven by sympathetic nerve activity and proteins secreted from BAT, to preserve energy homeostasis (Pereira *et al*, 2021; Piao *et al*, 2018; Schulz *et al*, 2013; Verdeguer *et al*, 2016). One way BAT function may be impaired is through disruption to mitochondrial function, such as that seen in brown adipocyte-specific Lkb1 or mitochondrial transcription factor A knockout rodent models (Masand *et al*, 2018). In these examples, there is a paradoxical increase in metabolic efficiency with mitochondrial dysfunction, even under high-fat diet conditions and irrespective of environmental temperature. This phenomenon implies a decoupling of the thermogenic activity of BAT from energy expenditure, which may be related to targeted suppression of mitochondrial DNA (mtDNA) gene expression and resulting electron transport chain (ETC) imbalances (Masand *et al*., 2018). Such imbalances may elicit a mitochondrial unfolded protein response (UPR^mt^) in BAT, which subsequently influences systemic metabolism independent of thermogenesis by promoting the appearance of beige adipocytes to compensate for BAT dysfunction and maintain metabolic stability. However, the precise interplay between BAT and WAT depots in response to the UPR^mt^ in BAT is unclear.

The mammalian UPR^mt^ involves a cascade of transcription factors, such as activating transcription factor 4 (ATF4), which are involved in optimizing thermoregulatory functions in BAT (Paulo *et al*, 2021). In particular, cold exposure upregulates ATF protein levels, emphasizing their integral role in maintaining thermal homeostasis. These transcription factors, including ATF4, ATF5, and C/EBP homologous protein (CHOP), promote mitochondrial biogenesis and expression of genes involved in protein folding quality control, and also mediate the release of mitokines like growth differentiation factor 15 (GDF15) and fibroblast growth factor 21 (FGF21), in response to the UPR^mt^ (Choi *et al*, 2020; Chung *et al*, 2017; Kang *et al*, 2021). However, the role of these mitokines in BAT where the UPR^mt^ is activated is not fully understood. The glial-derived neurotrophic factor (GDNF)-family receptor alpha-like (GFRAL) has been identified by four research groups as the receptor for GDF15, which signals through the coreceptor RET, primarily located in the hindbrain (Emmerson *et al*, 2017; Hsu *et al*, 2017; Mullican *et al*, 2017; Yang *et al*, 2017).

Our study reveals the pivotal role of GFRAL neurons in this context: we demonstrate that GDF15, secreted by brown adipocytes undergoing UPR^mt^, initiates beige adipogenesis through the activation of GFRAL neurons. This discovery underscores previously unrecognized BAT–brain crosstalk via the GDF15–GFRAL axis, which is activated by either physiological stress from cold exposure or pathological stress from mitochondrial dysfunction, and, in turn, orchestrates compensatory adaptive thermogenesis in beige adipocytes. Thus, GFRAL neurons emerge as master regulators in the thermogenic network, underpinning the relationship between BAT function and whole-body energy homeostasis.

## Results

### Induction of the UPR^mt^ and the integrated stress response in the BAT of Crif1 (CR6-interacting factor 1)-deficient mice

To investigate the effects of abnormalities in brown adipocyte-specific mitochondrial proteostasis, we generated a mouse model of mitochondrial dysfunction induced by the genetic inactivation of the *Crif1* gene in brown adipocytes. CRIF1 is critical to the function of the mitoribosome and the mitochondrial translation of mtDNA-encoded oxidative phosphorylation (OXPHOS) polypeptides. Inactivation of the *Crif1* gene results in significant reductions in the production of OXPHOS components and ETC activity (Kim *et al*, 2012). BAT-specific *Crif1*-knockout mice (Crif1^BKO^, generated using *Ucp1-Cre* and *Crif1^f/f^* mice) had significantly lower *Crif1* mRNA and protein expression in BAT than control mice (**Fig. 1A and Supplementary** Fig. 1A–C). In addition, Crif1^BKO^ mice had significantly lower expression of the OXPHOS complexes I (NDUFB8, nuclear DNA-encoded), III (UQCRC2, nuclear [n]DNA-encoded), and IV (MTCO1, mtDNA-encoded) than control mice (**Fig. 1A and Supplementary** Fig. 1C). Electron microscopy of brown adipocytes from Crif1^BKO^ mice revealed swollen mitochondria with abnormal cristae (**Fig. 1B**). Thus, the ablation of the *Crif1* gene in BAT results in substantial suppression of OXPHOS complex expression and mitochondrial structural abnormalities. The BAT of Crif1^BKO^ mice had larger lipid droplets and reduced multilocularity, indicative of potential defects in fatty acid oxidation and/or triglyceride storage (**Fig. 1C and Supplementary** Fig. 1D). UCP1 protein levels were either unchanged (males) or significantly increased (females) (**Fig. 1C and Supplementary** Fig. 1C).

**Figure 1.**
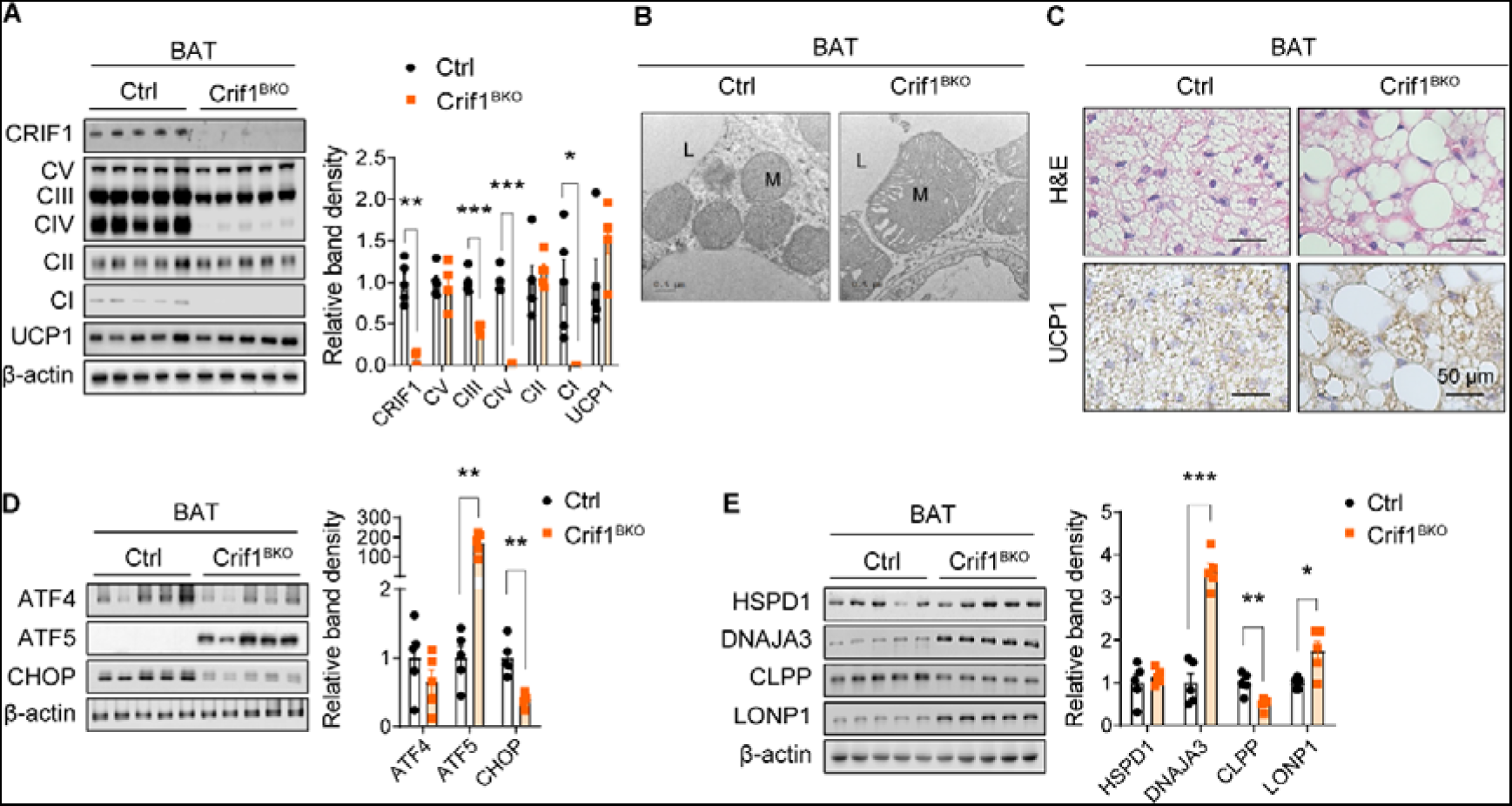
Crif1^BKO^ mice with a brown adipocyte-specific defect in oxidative phosphorylation show higher brown adipose tissue (BAT) expression of mitochondrial unfolded protein response proteins. A, Immunoblots and band densities for CRIF1, oxidative phosphorylation proteins, and UCP1 in the BAT of male control (Ctrl) and Crif1^BKO^ mice. B, Representative electron microscopic images of brown adipocytes. L: lipid, M: mitochondria. Bar, 0.5 μm. C, Hematoxylin and eosin (H&E)-stained and UCP1-immunostained sections of BAT. Bar, 50 μm. D, Immunoblots and band densities for mitochondrial unfolded protein response-associated transcription factors. E, Immunoblots and band densities for mitochondrial chaperones and proteins. *, *P* < 0.05; **, *P* < 0.01; ***, *P* < 0.001 *vs*. Ctrl by Student’s t-test. Data are represented as mean ± standard error of mean (SEM). See also Supplementary Figure 1.

Abnormal expression of OXPHOS components in *Crif1*-deficient cells induces transcriptomic changes, including those relevant to the UPR^mt^, which consists of both cell-autonomous and non-cell-autonomous responses (Chung *et al*., 2017). The ATF family, including ATF4 and ATF5, and CHOP are implicated in the mammalian UPR^mt^ (Fiorese *et al*, 2016; Quirós *et al*, 2017). In the present study, we evaluated the induction of the integrated stress response (ISR) and the UPR^mt^ in the BAT of Crif1^BKO^ mice and found that the expression of ATF5, but not ATF4 or CHOP, was significantly higher in the BAT of *Crif1*^BKO^ mice than control mice, suggesting a role for brown adipocytes in the maintenance of UPR^mt^ activity during mitochondrial stress (**Fig. 1D and Supplementary** Fig. 1E). In addition, mitochondrial chaperones and intrinsic proteases, such as heat shock protein family D member 1 (HSPD1), DnaJ heat shock protein family member A3 (DNAJA3), and lon peptidase 1 (LONP1), were expressed at significantly higher levels in the BAT of Crif1^BKO^ mice (**Fig. 1E and Supplementary** Fig. 1F). However, the expression of another mitochondrial intrinsic protease, caseinolytic mitochondrial matrix peptidase proteolytic subunit (CLPP), was significantly lower than in control mice in both male and female Crif1^BKO^ mice (**Fig. 1E and Supplementary** Fig. 1F).

These results demonstrate the consequences of mitoribosomal dysfunction in brown adipocytes: less biogenesis of OXPHOS components, structural abnormalities of the mitochondria, and an induction of both the ISR and UPR^mt^, as evidenced by elevated expression of ATF5, mitochondrial chaperones such as DNAJA3, and intrinsic proteases including LONP1.

### Metabolic characteristics of thermogenic compensation by Crif1^BKO^ mice subjected to cold exposure or a high-fat diet (HFD)

Crif1^BKO^ mice housed at room temperature (RT) underwent assessments for body weight, daily food intake, and the weight of various organs including adipose tissue, liver, and skeletal muscle. Both male and female Crif1^BKO^ mice exhibited significantly lower body weight and daily food intake compared with control mice. Both of the epidydimal and inguinal WAT mass in Crif1^BKO^ mice were lower than those in control mice (**Figs. 2A–C and Supplementary** Figs. 2A–C). An analysis of covariance showed that Crif1^BKO^ mice exhibited significantly higher energy expenditure (kcal/day) independent of body weight during paired feeding (**Figs. 2D,E**).

**Figure 2.**
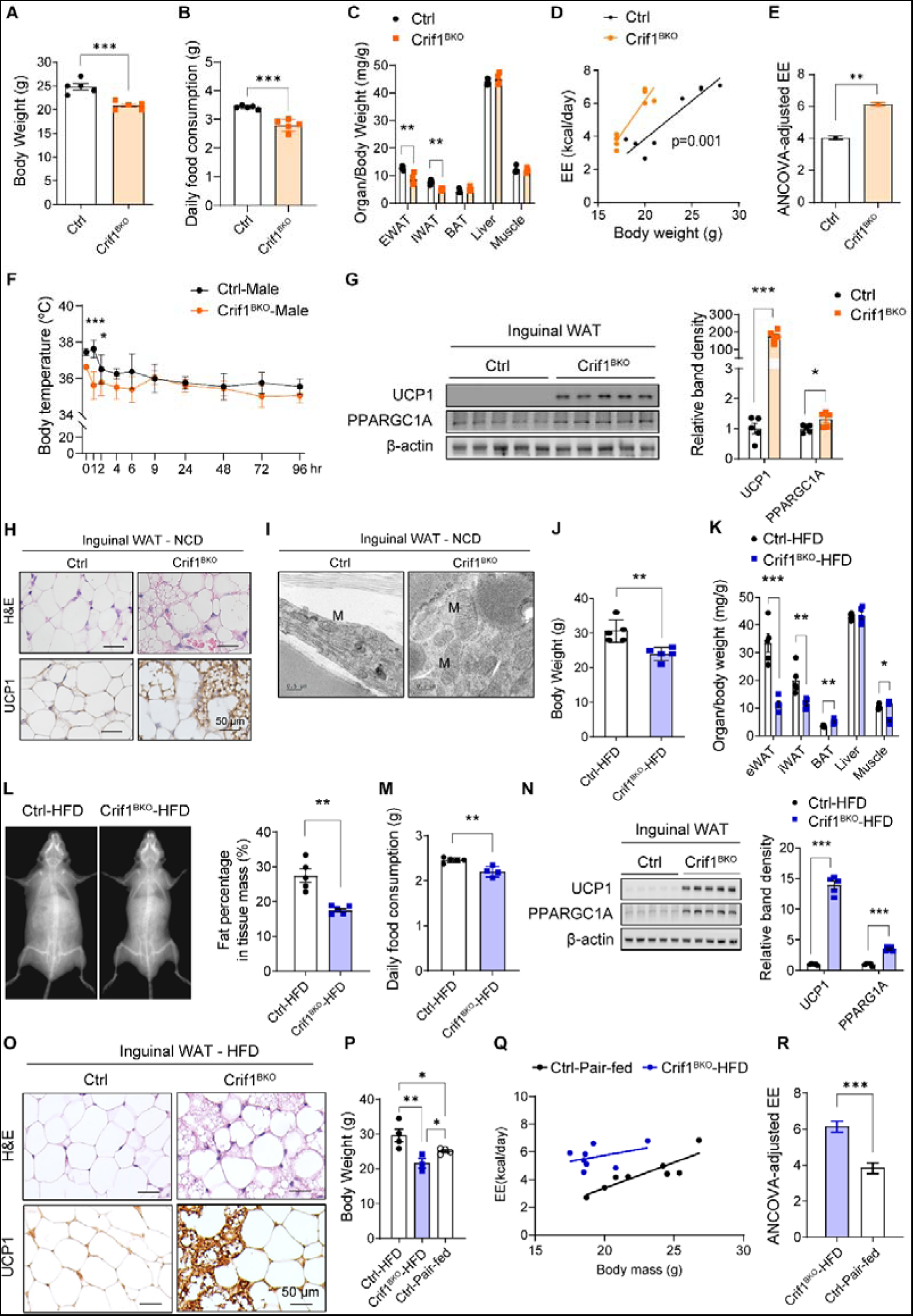
Crif1^BKO^ mice exhibit browning of inguinal white adipose tissue (WAT) and demonstrate resistance to high-fat diet-induced obesity. **A–C**, Body weight (**A**), daily food consumption (**B**), and organ/body weight ratios (**C**) for male control (Ctrl) and Crif1^BKO^ mice (n=5 per group). **D, E,** ANCOVA of energy expenditure (EE) against body weight (**D**) and ANCOVA-adjusted EE (**E**) in Ctrl and Crif1^BKO^ mice (n=8 per group) during paired feeding. **F**, Body temperatures of male Ctrl and Crif1^BKO^ mice housed at room temperature (22–24[) following cold exposure at 5[ for 1 week (n=4 per group). **G**, Immunoblots and band densities for UCP1 and PPARGC1A in inguinal WAT. **H**, Representative images of H&E-stained and UCP1-immunostained sections of inguinal WAT in mice fed a normal chow diet (NCD). Bar, 50 μm. **I,** Representative electron microscopic images of inguinal white adipocytes. L: lipid, M: mitochondria. Bar, 0.5 μm. **J–L**, Body weights (**J**), organ/body weight ratios (**K**), and fat percentages (**L**), according to densitometry. **M**, Daily food intake of male Ctrl and Crif1^BKO^ mice fed a HFD for 8 weeks (n=3–4 per group). **N**, Immunoblots and band densities for UCP1 and PPARGC1A in inguinal WAT. **O**, Representative images of H&E-stained and UCP1-immunostained sections of inguinal WAT. **P**, Body weight measured at 24 weeks of age after 18 weeks of HFD feeding in Ctrl (n=4), Crif1^BKO^ (n=3), and pair-fed Ctrl groups (n=4). **Q-R,** Energy expenditure of Crif1^BKO^ (n=8) and pair-fed Ctrl mice (n=8) after 18 weeks of HFD feeding. Bar, 50 μm. *, *P* < 0.05; **, *P* < 0.01; ***, *P* < 0.001 *vs*. Ctrl by Student’s t-test. Data are represented as mean ± SEM. See also Supplementary Figure 2 for *ad libitum* feeding results and Supplementary Figure 3 for HFD results.

To evaluate the ability of Crif1^BKO^ mice to maintain their body temperature in a cold environment, we measured body temperature after housing them at 5°C for 1 week. The body temperatures of Crif1^BKO^ mice were significantly lower than those of the control mice during the first 4 h of cold exposure (*P*<0.05), but they subsequently maintained body temperatures comparable with the control mice and showed no mortality (**Fig. 2F**). There were no differences in the glucose or insulin tolerance test results of control and Crif1^BKO^ mice (**Supplementary** Figs. 2D–G). These observations indicate that despite impaired BAT OXPHOS, Crif1^BKO^ mice have higher energy expenditure than control mice, although glucose metabolism was not affected.

Next, to evaluate the contribution of inguinal WAT to thermogenic compensation and systemic metabolic homeostasis, we measured the expression of UCP1 in this thermogenic fat depot of Crif1^BKO^ mice and found significantly higher expression of UCP1 and peroxisome proliferator-activated receptor gamma co-activator 1-alpha (PPARGC1A) than in control mice (**Fig. 2G and Supplementary** Fig. 2H). Histological examination revealed multilocular adipocytes and more UCP1 immunostaining in Crif1^BKO^ mice than in control mice (**Fig. 2H and Supplementary** Fig. 2I). Transmission electron microscopy revealed more abundant and larger mitochondria in the inguinal WAT of Crif1^BKO^ than in control mice (**Fig. 2I**). These findings suggest that WAT browning may contribute to the maintenance of systemic energy homeostasis despite BAT OXPHOS defects in Crif1^BKO^ mice.

To further characterize the differences in the systemic metabolism of Crif1^BKO^ mice, we evaluated metabolic parameters after HFD feeding for 12 weeks. Crif1^BKO^ mice fed an HFD (Crif1^BKO^-HFD) had significantly lower expression of CRIF1; the OXPHOS complexes I, III, and IV; and UCP1 (**Supplementary** Figs. 3A–C). Crif1^BKO^ mice fed an HFD also had significantly higher expression of DNAJA3 and LONP1 in BAT, although CLPP was expressed at a significantly lower level (**Supplementary** Fig. 3A). The body weight of Crif1^BKO^-HFD mice was significantly lower than that of control mice (**Fig. 2J and Supplementary** Fig. 3C), and the WAT mass and fat percentages, determined using dual-energy X-ray absorptiometry, were significantly lower in Crif1^BKO^ mice, at only one-third or one-half of those of the control group (**Figs. 2K,L and Supplementary** Figs. 3D,E). The daily food consumption of male Crif1^BKO^-HFD mice was lower than that of the controls (**Fig. 2M and Supplementary** Fig. 3F). Glucose and insulin tolerance testing showed no differences between the control and Crif1^BKO^-HFD mice (**Supplementary** Figs. 3G–J). The inguinal WAT of Crif1^BKO^-HFD mice had significantly higher expression of UCP1 and PPARGC1A, a greater number of multilocular adipocytes, and greater UCP1 immunostaining (**Figs. 2N,O and Supplementary** Figs. 3K,L), consistent with the results obtained in normal chow-fed Crif1^BKO^ mice (**Figs. 2G,H and Supplementary** Figs. 2H,I). Furthermore, the body weight of Crif1^BKO^ mice exhibited significantly decrease compared to pair-fed control after 18 weeks of high-fat diet, indicating that Crif1^BKO^ mice had a change of energy expenditure in addition to decreased food intake (**Fig 2P**). As anticipated, energy expenditure (kcal/day) of Crif1^BKO^-HFD mice was significantly higher after adjusting for body weight through analysis of covariance compared with pair-fed control mice (**Fig 2Q,R**). In summary, Crif1^BKO^ mice exhibited resistance to weight gain when fed an HFD. In addition, Crif1^BKO^ mice consumed less food, had higher energy expenditure, and had more beige adipocytes in the inguinal WAT depot than control mice.

Taken together, these findings suggest that Crif1^BKO^ mice maintain a lower body weight and higher energy expenditure than control mice, despite impaired BAT OXPHOS. Moreover, when fed an HFD, Crif1^BKO^ mice resist weight gain, effects associated with increased browning of the inguinal WAT depot.

### Identification and mechanism of GDF15 expression by the UPR^mt^ in the BAT of Crif1^BKO^ mice

To confirm the non-cell-autonomous effect of UPR^mt^ activation in BAT, we aimed to identify proteins secreted from BAT using both the UniProt database (www.uniprot.org) and RNA-seq data. Bulk RNA-seq of Crif1^BKO^ BAT showed 54 genes encoding secretory proteins that were significantly differentially expressed, with log_2_-fold differences >2 and adjusted *P*-values <0.01 (**Fig. 3A**). Of these, the expression of *Gdf15* had one of the highest fold changes and lowest *P*-values. This is consistent with previous findings that GDF15 is a mitokine, which in mice and humans is secreted during mitochondrial dysfunction (Choi *et al*., 2020; Chung *et al*., 2017; Kang *et al*., 2021; Khan *et al*, 2017; Montero *et al*, 2016). The difference in BAT gene expression was accompanied by significantly higher plasma concentrations of GDF15 in Crif1^BKO^ mice fed either normal chow or an HFD than control mice (**Figs. 3B–E and Supplementary** Figs. 4A–D). By contrast, the expression of *Fgf21* and serum levels of FGF21 in Crif1^BKO^ mice were significantly higher than control mice only with a normal chow diet, with no significant changes observed under HFD feeding (**Supplementary** Figs. 4E–L). After both a normal chow diet and HFD feeding, *Gdf15* and *Fgf21* mRNA levels in the liver were not significantly different between control and Crif1^BKO^ mice, except for *Fgf21* expression under normal chow feeding in females (**Supplementary** Figs. 4M–P).

**Figure 3.**
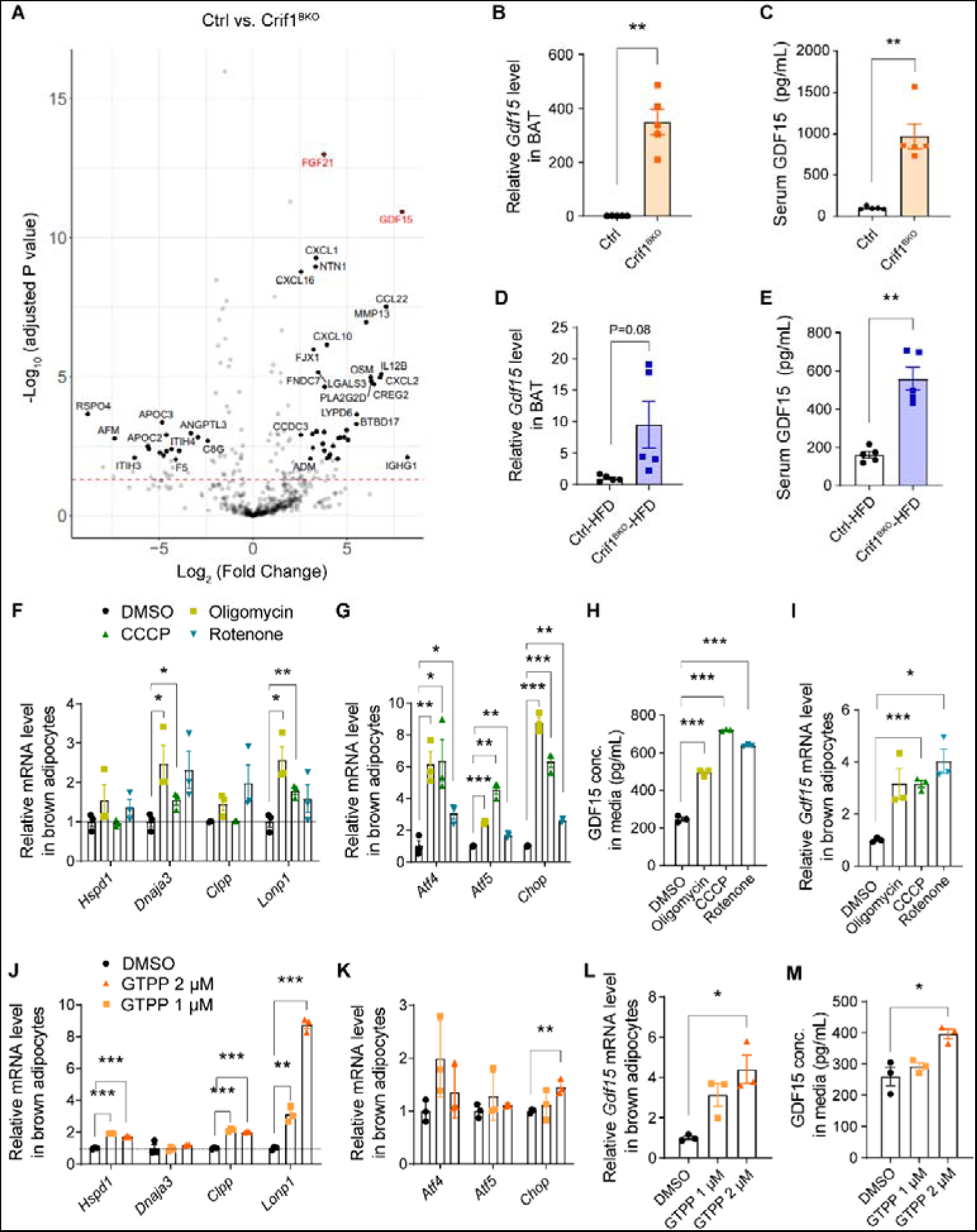
*Gdf15* expression increases in response to mitochondrial unfolded protein response activation in brown adipose tissue (BAT). **A**, Volcano plot of differentially expressed secretory protein-encoding genes in the BAT of Crif1^BKO^ mice, identified using RNA sequencing. **B**, Relative *Gdf15* expression in the BAT of male control (Ctrl) and Crif1^BKO^ mice (n=5 per group), as characterized in Fig. 1. **C**, Serum GDF15 concentrations in male Ctrl and Crif1^BKO^ mice. **D**, Relative *Gdf15* expression in the BAT of the male Ctrl-HFD and Crif1^BKO^-HFD mice, as characterized in Fig. 3. **E**, Serum GDF15 concentration in male Ctrl-HFD and Crif1^BKO^-HFD mice. **F–I**, Relative mRNA expression of mitochondrial unfolded protein response-associated genes (**F**), transcription factors (**G**), and *Gdf15* (**H**) in differentiated brown adipocytes treated with oxidative phosphorylation inhibitors (10 μg/mL oligomycin, 2 μg/mL CCCP, or 1 μM rotenone) for 24 h. (**I**) GDF15 concentration in the medium. **J–L,** Relative mRNA expression of mitochondrial unfolded protein response-associated genes (**J**), transcription factors (**K**), and *Gdf15* (**L**) in differentiated brown adipocytes treated with gamitrinib (G-TTP) for 24 h. **M**, GDF15 concentration in cell medium. *, *P* < 0.05; **, *P* < 0.01; ***, *P* < 0.001 *vs*. 0 h, Ctrl, Ctrl-HFD, or DMSO by Student’s t-test. Data are represented as mean ± SEM. See also Supplementary Figure 4.

To further characterize the relationship between mitochondrial OXPHOS dysfunction, UPR^mt^ activation, and GDF15 secretion, we treated differentiated immortalized brown adipocytes with OXPHOS complex inhibitors and measured the expression of genes encoding proteins involved in the ISR and UPR^mt^ pathways. The expression of the mitochondrial chaperone *Dnaja3* and the intrinsic protease *Lonp1* was significantly higher after treatment with 10 μg/mL oligomycin (a complex V inhibitor) and 2 μg/mL carbonyl cyanide m-chlorphenyl hydrazine (CCCP, an uncoupler), and expression of the transcription factors *Atf4*, *Atf5*, and *Chop* was significantly higher after treatment with oligomycin, CCCP, and 1 μM rotenone (a complex I inhibitor) treatment than after control treatment (DMSO) (**Figs. 3F,G**). In addition, brown adipocytes treated with OXPHOS complex inhibitors had significantly higher *Gdf15* mRNA expression and a 2–3-fold higher concentration of GDF15 in the medium, with similar responses seen in the mitokine FGF21 (**Figs. 3H,I and Supplementary** Figs. 4Q,R). To investigate the connection between UPR^mt^ and the induction of mitokine expression in brown adipocytes, we induced cellular proteostasis stress in the mitochondrial matrix using gamitrinib-triphenylphosphonium (GTPP), an Hsp90 inhibitor (Münch & Harper, 2016). As expected, GTPP treatment resulted in significantly higher mRNA expression of *Hspd1*, *Clpp*, and *Lonp1* than after the control treatment (**Fig. 3J**). The expression of *Chop* and *Gdf15* and the concentration of GDF15 in the medium was also significantly higher after 2 μM GTPP treatment than after control treatment (**Figs. 3K – M**). In addition, the concentration of FGF21 in the medium was significantly higher after 2 μM GTPP treatment than after control treatment, in line with the tendency toward higher expression observed at the mRNA level (**Supplementary** Figs. 4S,T).

Overall, these results indicate that GDF15 production increases in BAT in response to OXPHOS dysfunction and abnormalities in proteostasis, induced by both pharmacological intervention and genetic manipulation, *in vitro* and *in vivo*. The higher resulting concentrations of GDF15 in Crif1^BKO^ mice likely affect systemic metabolism and adaptive thermogenesis.

### Induction of the UPR^mt^ and GDF15 expression in the BAT of mice during cold exposure

The above results demonstrate that in Crif1^BKO^ mice the UPR^mt^ is induced, proteostasis is dysfunctional, and GDF15 expression is increased, which is compensated for by adaptive thermogenesis in inguinal WAT. To determine the effects of environmental, rather than genetic, BAT manipulation, we assessed UPR^mt^ activation and GDF15 expression in mice exposed to cold. Cold exposure increases mitochondrial demand in thermogenic adipose tissue and triggers induction of the mitochondrial stress response (Chouchani *et al*, 2016; Forner *et al*, 2009; Lu *et al*, 2018). We assessed the induction of the UPR^mt^ in the BAT of C57BL/6 wild-type (WT) mice in response to cold exposure (7°C) for 24 h. After 6 h of exposure, the expression of *Ucp1* mRNA in BAT was 26-fold higher than at baseline (**Fig. 4A**). We measured the expression of genes involved in the UPR^mt^ and found that expression of the mitochondrial chaperones *Hspd1* and *Dnaja3*, the mitochondrial matrix proteases *Clpp* and *Lonp1*, and the inner-membrane protease *Yme1ll* was 2–5-fold higher after 6 and 24 h of cold exposure than at baseline (**Figs. 4B–E**), but expression of the inner-membrane protease *Imm2l* and the outer-membrane protease *Usp30* was not affected. The mRNA expression of *Atf4* and *Atf5*, transcription factors known to regulate the UPR^mt^, was approximately 20-fold higher after 6 and 24 h of cold exposure than at baseline (**Fig. 4F**), and the protein expression of ATF4 and CHOP, but not that of ATF5, was also significantly higher after cold exposure (**Fig. 4G**). Thus, the expression of mitochondrial chaperones and proteases involved in the UPR^mt^ was increased in the BAT of mice during cold-induced thermogenesis. In addition, the expression of the mitokines *Gdf15* and *Fgf21* was significantly higher in the BAT of cold-exposed mice after 6 and 24 h of cold exposure than at baseline, as were the serum concentrations of GDF15 and FGF21 after 24 h of cold exposure (**Figs. 4H,I and Supplementary** Figs. 5A,B).

**Figure 4.**
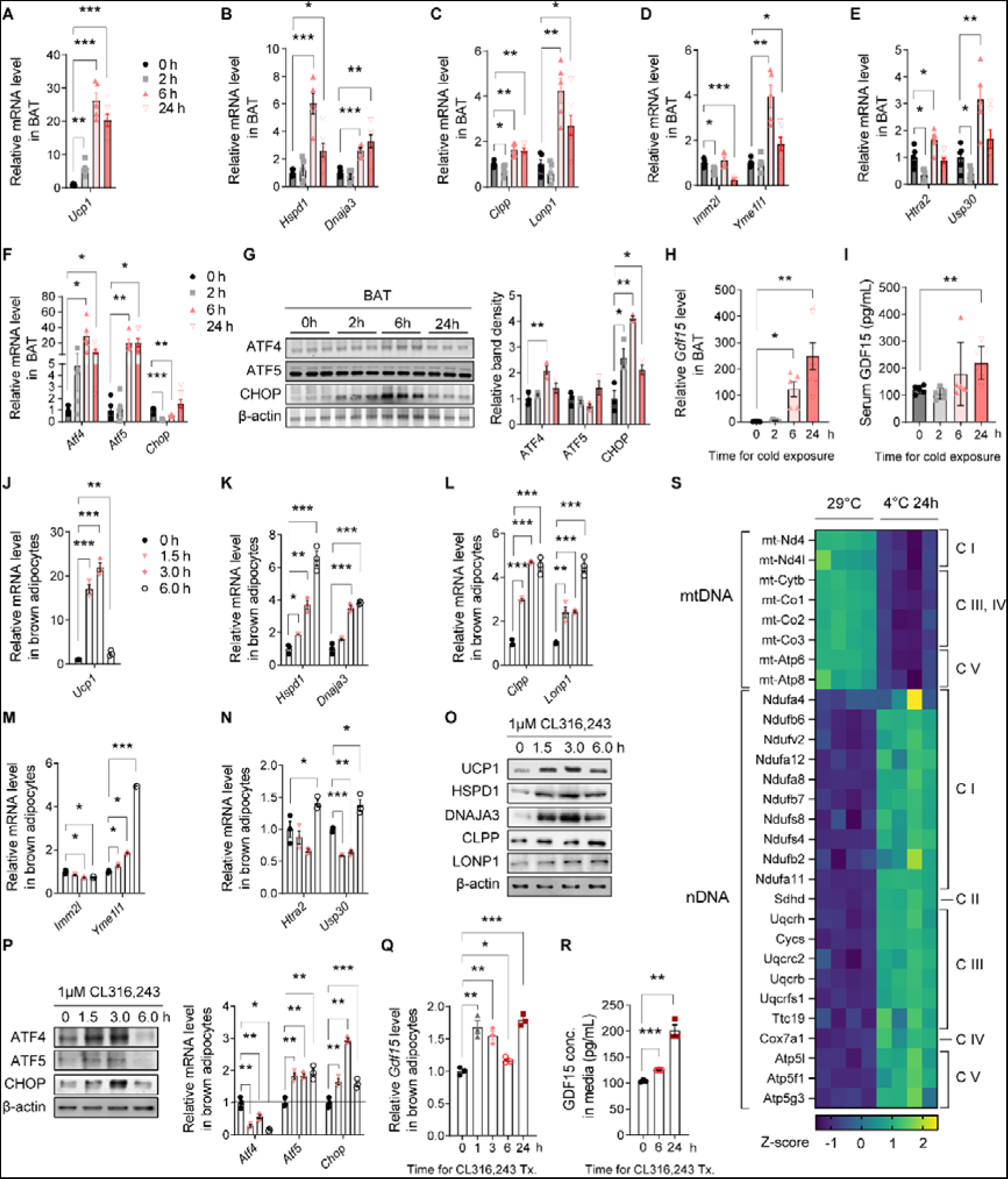
The expression of mitochondrial chaperones and proteases is high in the brown adipose tissue (BAT) of mice with cold-induced thermogenesis. **A–F**, Relative mRNA expression of *Ucp1* (**A**), and genes encoding mitochondrial chaperones (**B**), mitochondrial matrix proteins (**C**), inner-membrane (**D**), inter- and outer-membrane proteases (**E**), and transcription factors (**F**) in BAT from wild-type mice housed at 30°C following cold exposure at 5°C for 24 h (n=5–7 per group). **G**, Representative immunoblots and band densities for transcription factors in the BAT of cold-exposed mice. **H**, Relative GDF15 expression in the BAT of cold-exposed mice. (**I**) Serum GDF15 concentrations of cold-exposed mice. **J–N**, Relative mRNA expression of *Ucp1* (**J**), mitochondrial chaperones (**K**) and proteases **L-N,** in differentiated immortalized brown adipocytes treated with 1 µM CL 316,243. **O,** Representative immunoblots for UCP1, mitochondrial chaperones, and proteases. **P**, Representative immunoblots for transcription factors and relative mRNA expression. **Q**,**R**, Relative *Gdf15* expression and GDF15 concentration in cell medium. **s**, Heatmap of differentially expressed oxidative phosphorylation genes encoded by mitochondrial DNA (mtDNA) and nuclear DNA (nDNA) from previously published RNA-seq data^26^. *, *P* < 0.05; **, *P* < 0.01; ***, *P* < 0.001 *vs*. 0 h by Student’s t-test. Data are represented as mean ± SEM. See also Supplementary Figure 5.

To further explore the induction of UPR^mt^ *in vitro*, we treated differentiated immortalized brown adipocytes with 1 µM CL 316,243, a β3-agonist. *Ucp1* expression in these cells was ∼20-fold higher after 3 h of treatment than at baseline (**Fig. 4J**), and the mRNA expression of mitochondrial chaperones (*Hspd1* and *Dnaja3*) and proteases (*Clpp*, *Lonp1*, *Yme1l1*, *Htra2*, and *Usp30*) was 4–6-fold higher after 6 h of treatment, while *Imm2l* expression was significantly lower after 3 and 6 h of treatment than at baseline (**Figs. 4K–N**). The protein expression of HSPD1, DNAJA3, CLPP, and LONP1 was also higher after CL 316,243 treatment than at baseline (**Fig. 4O**). The expression of the related transcription factors *Atf5* and *Chop* was significantly higher, while expression of *Atf4* was significantly lower, from 1.5 to 3 h of treatment compared with baseline (**Fig. 4P**). The mRNA expression of *Gdf15* and *Fgf21*, as well as the concentration of GDF15 and FGF21 in the media, was also significantly higher in differentiated immortalized brown adipocytes after CL 316,243 treatment than at baseline (**Figs. 4Q,R and Supplementary** Figs. 5C,D). Additionally, the expression of *Ucp1* and mitochondrial chaperones and proteases in undifferentiated brown adipocytes were only slightly higher (2–4-fold higher than controls) after 1.5 h of treatment with 1 µM CL 316,243 (**Supplementary** Figs. 5E–I). The mRNA expression of *Gdf15*, *Fgf21*, *Atf4*, *Atf5*, and *Chop* was significantly lower in undifferentiated brown adipocytes, unlike the differentiated cells, which have the mitochondrial capacity for thermogenesis (**Supplementary** Figs. 5J,K). These results demonstrate a specific induction of the UPR^mt^ and mitokine secretion in thermogenic brown adipocytes, as evidenced by greater induction of the UPR^mt^ in differentiated cells than in undifferentiated cells.

Previous studies show that an imbalance between nDNA and mtDNA-encoded OXPHOS proteins activates the UPR^mt^ (Houtkooper *et al*, 2013). To determine whether this mitochondrial/nuclear protein imbalance was present in cold-exposed mice, we measured the mRNA expression of OXPHOS components in previously published RNA sequencing data from the BAT of C57BL/6 mice after 24 h of cold exposure (4°C) (Quesada-López *et al*, 2016). Of the 13 mtDNA-encoded OXPHOS complex genes, eight were present at significantly lower levels after cold exposure, and of the nDNA-encoded oxidative phosphorylation proteins, 21 were present at significantly higher levels (**Fig. 4S**). We therefore concluded that these changes in transcript levels would induce a mitochondrial/nuclear protein imbalance that may have been responsible for the induction of the UPR^mt^ in cold-exposed BAT.

In summary, cold exposure increases mitochondrial demand in BAT, resulting in UPR^mt^ induction, both in mice subjected to cold exposure and in differentiated immortalized brown adipocytes treated with a β3-agonist. *Ucp1*, mitochondrial chaperones and proteases, and mitokines increase in cold-exposed BAT and *in vitro* in β3-agonist-exposed differentiated brown adipocytes, but less so in undifferentiated brown adipocytes. Activation of the UPR^mt^ in cold-exposed mice may result from an imbalance between nDNA and mtDNA-encoded OXPHOS proteins.

### The UPR^mt^ and Gdf15 expression in the BAT of mice is low under thermoneutral conditions

As shown in Fig. 4, the UPR^mt^ was induced by cold exposure; however, RT (22– 24°C) housing of mice also causes thermal stress (Feldmann *et al*, 2009). To avoid thermal stress in Crif1^BKO^ mice, animals were housed under thermoneutral conditions (30°C) for 8 weeks, which resulted in lower UPR^mt^-related protein and mRNA expression, as well as significantly lower UCP1 expression, in the BAT of 30°C-housed Crif1^BKO^ mice than RT-housed Crif1^BKO^ mice (**Figs. 5A,B**). At both RT and 30°C, Crif1^BKO^ mice had lower protein expression of OXPHOS complexes I, III, and IV, as well as lower *Crif1* mRNA expression, than control mice housed in the same conditions (**Supplementary** Figs. 6A,B). On histological examination, both Crif1^BKO^ and control mice were found to have unilocular adipocytes and low UCP1 expression under thermoneutral conditions (**Supplementary** Fig. 6C). mRNA expression of *Gdf15* in BAT and the concentration of serum GDF15 in the 30°C-housed Crif1^BKO^ mice was also significantly lower than RT-housed Crif1^BKO^ mice, implying an attenuated UPR^mt^ response (**Figs. 5C,D**). mRNA expression of *Fgf21* in the BAT of 30°C-housed Crif1^BKO^ mice was significantly lower than RT-housed Crif1^BKO^ mice, while serum FGF21 concentration was not different between Ctrl and Crif1^BKO^ mice in 30°C housing (**Supplementary** Figs. 6D,E).

**Figure 5.**
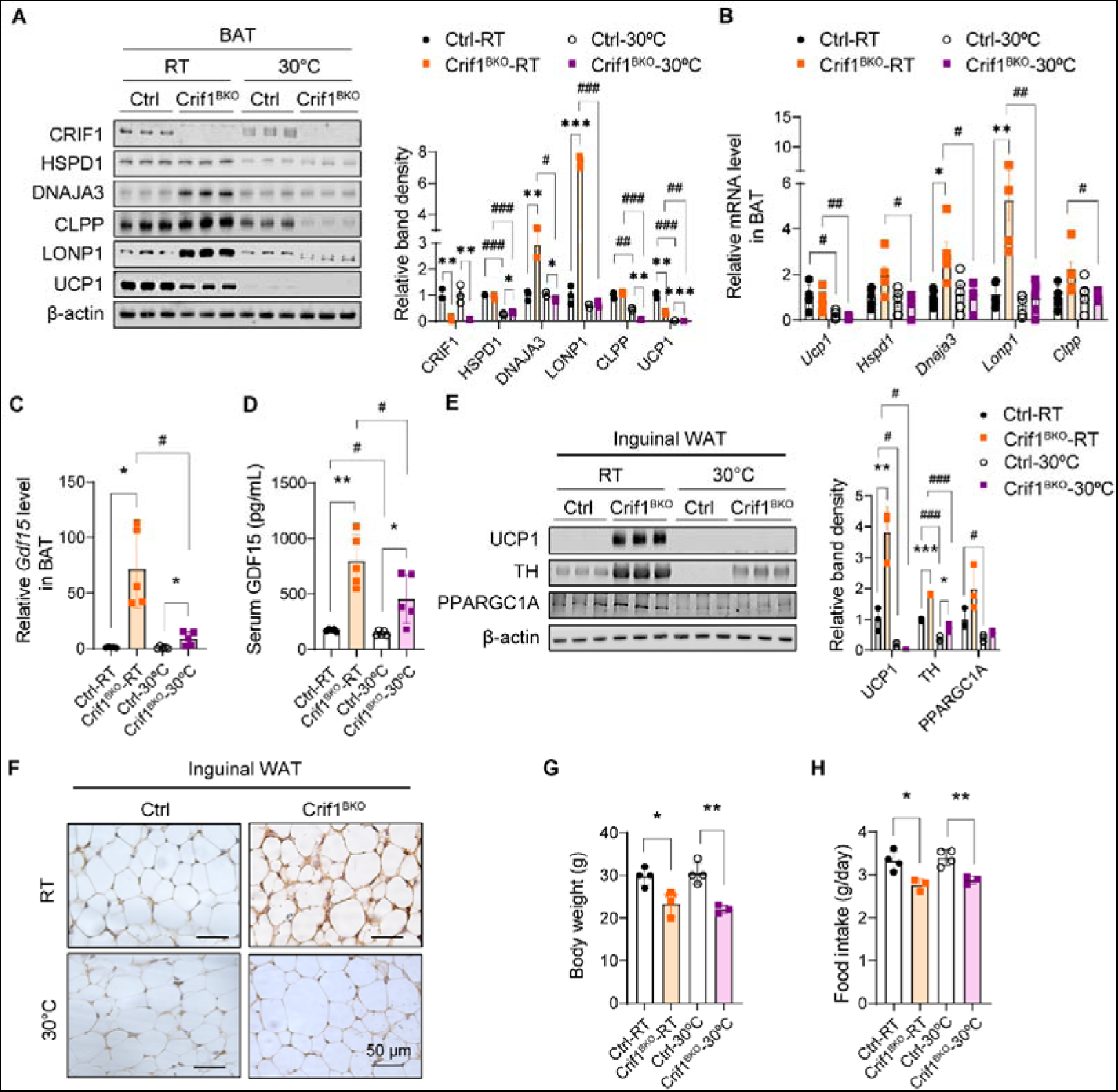
Activation of the mitochondrial unfolded protein response and *Gdf15* expression in brown adipose tissue (BAT) is low under thermoneutral conditions. A, Immunoblots and band densities for CRIF1, mitochondrial unfolded protein response, and UCP1 in the BAT of 12–13-week-old male control (Ctrl) and Crif1^BKO^ mice housed at room temperature (22–24°C) (Ctrl-RT or Crif1^BKO^-RT) or 30°C (Ctrl-30°C or Crif1^BKO^-30°C) for 8 weeks. **B**, Relative mRNA expression of *Ucp1* and mitochondrial unfolded protein response genes in BAT. **C**, Relative *Gdf15* expression in BAT. **D**, Serum GDF15 concentration. **E**, Immunoblots and band densities for UCP1, tyrosine hydroxylase (TH), and PPARGC1A in inguinal white adipose tissue (WAT). **F**, Representative images of UCP1-immunostained inguinal WAT. Bar, 50 μm. **G**,**H**, Body weight (**G**) and daily food intake (**H**) of 15–16-week-old male Ctrl and Crif1^BKO^ mice housed at room temperature (22–24°C) (Ctrl-RT or Crif1^BKO^-RT) or 30°C (Ctrl-30°C or Crif1^BKO^-30°C) for 9 weeks (n=3–4 per group). *, *P* < 0.05; **, *P* < 0.01; ***, *P* < 0.001 *vs*. Ctrl by Student’s t-test; #, *P* < 0.05; ##, *P* < 0.01; ###, *P* < 0.001 *vs.* RT by Student’s t-test. Data are represented as mean ± SEM. See also Supplementary Figure 6.

In the inguinal WAT of Crif1^BKO^ mice, the expression of UCP1 and tyrosine hydroxylase (TH), the rate-limiting enzyme in catecholamine synthesis, was significantly higher in RT-housed mice than 30°C-housed mice (**Figs. 5E,F**). *Gdf15* expression in the livers of Crif1^BKO^ mice did not differ from that of control mice (**Supplementary** Fig. 6F). The body weights of Crif1^BKO^ mice were significantly lower than control mice (**Fig. 5G and Supplementary** Figs. 6G,H), which is consistent with significantly higher serum GDF15 concentrations (**Fig. 5D**) and significantly lower food intakes of 30°C-housed Crif1^BKO^ mice than control mice housed at the same temperature (**Fig. 5H**).

These results suggest that under thermoneutral conditions, an absence of thermal stress blunts the UPR^mt^ activation, as well as elevated *Gdf15* mRNA expression and serum GDF15 concentrations observed in Crif1^BKO^ mice. Under these conditions, adaptive thermogenesis in the BAT of Crif1^BKO^ mice is not required, meaning mitochondrial proteostasis continues to function, even with OXPHOS deficits. Therefore, sub-thermoneutral stress amplifies UPR^mt^ activation in BAT, triggering a response in inguinal WAT via GDF15 expression.

### High circulating concentrations of GDF15 in Crif1^BKO^ mice activate GFRAL-positive neurons in the hindbrain

Neurons located in the area postrema (AP) and the nucleus of the solitary tract (NTS) regions of the hindbrain are well positioned to detect blood-borne cues and project neurites into the perivascular spaces of local fenestrated capillaries. Therefore, the AP is involved in interoception and is a key detector of several peptides, including GDF15 and GLP-1, which play important roles in the regulation of feeding behavior and metabolism (Tsai *et al*, 2019; Zhang *et al*, 2021). Key studies demonstrate that GFRAL is the central mediator of the metabolic effects of GDF15 (Emmerson *et al*., 2017; Hsu *et al*., 2017; Mullican *et al*., 2017; Yang *et al*., 2017). These studies converge on the AP and NTS in the brainstem as the primary sites of GFRAL expression, linking GDF15 signaling to reduced food intake and body weight regulation in rodent and nonhuman primate models (Emmerson *et al*., 2017; Hsu *et al*., 2017; Mullican *et al*., 2017; Yang *et al*., 2017).

To determine whether high circulating concentrations of GDF15 in Crif1^BKO^ mice activate subsets of glutamatergic neurons, including GFRAL-positive neurons, we assessed c-Fos expression in GFRAL-positive neurons in the hindbrain of control and Crif1^BKO^ mice. We first examined the distribution of GFRAL-positive neurons in the AP and NTS of WT mice using immunohistochemical staining. GFRAL-positive neurons were concentrated in the AP and NTS (**Fig. 6A**). By contrast, the paraventricular nucleus of the hypothalamus (PVH) and arcuate nucleus, which regulate metabolism, did not have GFRAL immunostaining (**Figs. 6B,C**). Next, we sectioned the brains of Crif1^BKO^ and control mice at the level of the medulla oblongata, focusing on the AP and NTS, and performed GFRAL and c-Fos immunostaining. We found that the c-Fos expression of GFRAL-stained cells was higher in Crif1^BKO^ mice than in controls, especially in the AP (**Figs. 6D,E**). This implies that GFRAL activity is higher in the AP of Crif1^BKO^ mice than controls, which may account for the lower food consumption of Crif1^BKO^-HFD mice (**Fig. 2M**).

**Figure 6.**
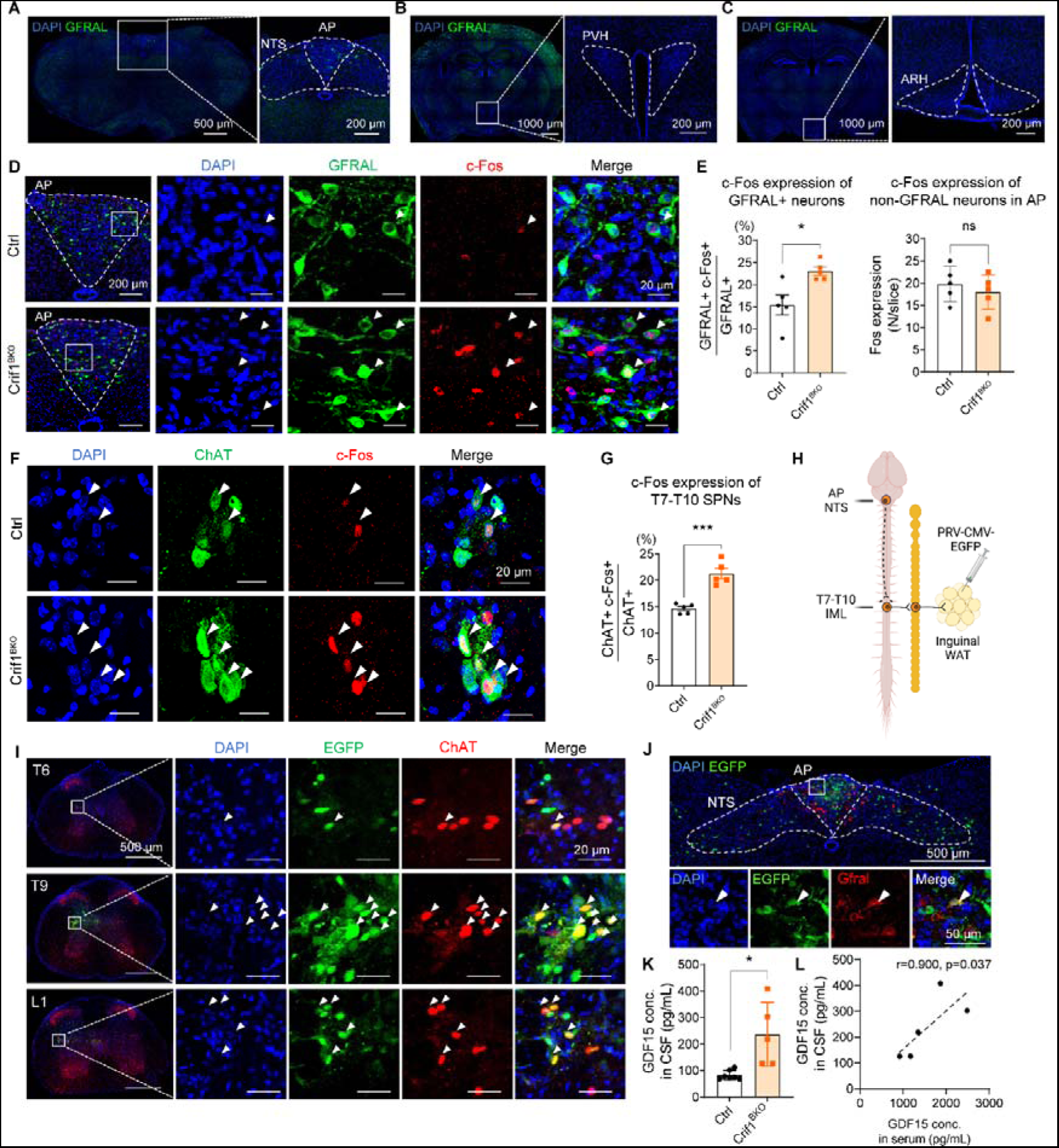
Activation of GFRAL-positive neurons in the hindbrain, combined with retrograde neuronal tracing of both the sympathetic nervous system and GFRAL neurons in Crif1^BKO^ mice. **A–C**, Representative images of the area postrema (AP) and nucleus of the solitary tract (NTS) (**A**), paraventricular nucleus of the hypothalamus (PVH; **B**), and arcuate nucleus (ARH; **C**) of wild-type mice stained with DAPI and immunostained for GFRAL. **D**, Representative images of the AP and NTS of control (Ctrl) and Crif1^BKO^ mice stained using DAPI and immunostained for GFRAL and c-Fos. **E**, c-Fos expression in GFRAL-expressing cells and non-GFRAL-expressing neurons in the AP and NTS from control mice (n=5) and Crif1^BKO^ mice (n=5). **F**, Representative images of the intermediolateral nucleus region of the thoracic spinal cords of Ctrl and Crif1^BKO^ mice stained with DAPI and immunostained for ChAT and c-Fos. **G**, c-Fos expression in ChAT-expressing cells. **H**, Schematic model of retrograde neuronal tracing study using a pseudorabies virus that express EGFP under CMV promoter (PRV-CMV-EGFP). AP, area postrema; NTS, nucleus tractus solitarius; IML, intermediolateral nucleus; WAT, white adipose tissue. **I**, Representative images of the intermediolateral nucleus region of the spinal cord of ChAT-IRES-Cre:tdTomato mice, 96 h after injection of enhanced green fluorescent protein (EGFP)-expressing pseudorabies virus into the inguinal WAT, stained with DAPI, and immunostained for EGFP and ChAT. **J**, Representative images of the AP and NTS of wild-type mice 120 h after injection of EGFP-expressing pseudorabies virus into the inguinal WAT, stained with DAPI and immunostained for EGFP and GFRAL. **K**, GDF15 concentrations in the cerebrospinal fluid of Ctrl and Crif1^BKO^ mice. **L**, Correlation between the serum and cerebrospinal fluid GDF15 concentrations in Crif1^BKO^ mice. *, *P* < 0.05; ***, *P* < 0.001 *vs*. Ctrl by Student’s t-test. Data are represented as mean ± SEM. See also Supplementary Figure 7.

Taken together, these results suggest that high circulating concentrations of GDF15 in Crif1^BKO^ mice are sufficient to activate GFRAL-positive neurons in the AP and NTS of the hindbrain, which may explain the metabolic phenotype and resistance to obesity of Crif1^BKO^ mice.

### Retrograde neuronal tracing reveals the role of the sympathetic nervous system and GFRAL-positive neurons in the GDF15-mediated browning of inguinal WAT in Crif1^BKO^ mice

We next investigated the involvement of sympathetic neuronal activation in the browning of inguinal WAT in Crif1^BKO^ mice (Huesing *et al*, 2021). c-Fos expression was significantly higher in the sympathetic preganglionic neurons of the intermediolateral nucleus region of the T7–T10 segment of the spinal cord, which innervates the inguinal WAT, of Crif1^BKO^ mice than control mice (**Figs. 6F,G**). To confirm communication between GFRAL-stained neurons in the brain and sympathetic preganglionic neurons of the spinal cord, we performed retrograde neuronal tracing using a pseudorabies virus. We injected an enhanced green fluorescent protein (EGFP)-expressing pseudorabies virus unilaterally into the inguinal WAT of ChAT-IRES-Cre:tdTomato mice, which express tdTomato in their sympathetic preganglionic neurons, and WT mice (**Fig 6H**). Ninety-six (96) h post-injection, EGFP-stained neurons were present in the intermediolateral nucleus region of the T6–L1 segment alone of the T1–L2 spinal cord and in multiple regions of the brain, including the paraventricular nucleus of the hypothalamus, locus coeruleus, and nucleus raphe pallidus (**Fig. 6I and Supplementary** Fig. 7A). At 120 h post-injection, EGFP-stained neurons were present in a wider range of brain regions, including the AP and NTS, primary motor cortex, hypothalamus, and reticular nuclei (**Supplementary** Fig. 7B). In the AP and NTS of the mice (n = 2), 11.8% and 27.8% of GFRAL-stained neurons colocalized with EGFP, respectively (**Fig. 6J and Supplementary** Fig. 7C). Moreover, compared with baseline levels, 24 h of cold exposure in control mice resulted in a pronounced increase in c-Fos expression in GFRAL-positive neurons (**Supplementary** Fig. 7D,E). These results suggest that GDF15-induced activation of GFRAL in the AP and NTS could activate sympathetic nerves innervating inguinal WAT and induce browning, as in Crif1^BKO^ and/or cold-induced mice (**Figs. 2G,H and Figs. 5E,F**).

We next investigated whether GDF15 activates the preganglionic sympathetic neurons of the spinal cord directly through cerebrospinal fluid (CSF). Significantly higher concentrations of GDF15 were found in the CSF of Crif1^BKO^ mice than control mice, and the CSF concentration was significantly positively correlated with the serum concentration (**Figs. 6K,L**). However, GFRAL was not detected by RNA fluorescence *in situ* hybridization in the spinal cords of WT C57BL/6 mice (**Supplementary** Figs. 7F). This suggests that the greater GFRAL activity in the AP of Crif1^BKO^ mice was caused by higher serum and/or CSF GDF15 concentrations activating the preganglionic sympathetic nerves of the spinal cord. Despite this activation, the blood pressure of Crif1^BKO^ mice was significantly lower, and their pulse rates did not differ from those of control mice (**Supplementary** Figs. 7G,H).

Taken together, these findings indicate that activation of GFRAL in the AP and NTS of Crif1^BKO^ mice by high serum and/or CSF GDF15 concentrations activates the preganglionic SNS in the spinal cord, leading to the browning of inguinal WAT. They suggest a direct link between systemic metabolism and defective adaptive thermogenesis mediated by GDF15 in mice with BAT mitochondrial dysfunction.

### *Gfral* antisense oligonucleotides prevent GDF15-mediated phenotypic changes in Crif1^BKO^ mice

To assess the role of GFRAL in adaptive thermogenesis within Crif1^BAT^, we synthesized antisense oligonucleotides (ASOs) targeting the *Gfral* gene. *Gfral* ASOs were designed to anneal to specific complementary regions of *Gfral* mRNA, triggering RNase H-mediated degradation (**Fig. 7A**). The phosphorothioate-modified backbone and bases enhance *Gfral* ASO stability and reduce immunogenicity, ensuring sustained knockdown (**Fig. 7B**). We explored *Gfral* suppression in the AP and NTS of the hindbrain using mouse models, with *Gfral* depletion achieved via intracerebroventricular (ICV) injections of *Gfral* ASOs. Crif1^BKO^ mice (n = 5 per group) fed *ad libitum* over 12 weeks received either *Gfral* ASOs or control ASOs in the final 2 weeks. RNA scope analysis of the hindbrain showed a significant 90% reduction in *Gfral* mRNA in the AP after 2 weeks of treatment with *Gfral* ASOs (**Figs. 7C–E**).

**Figure 7.**
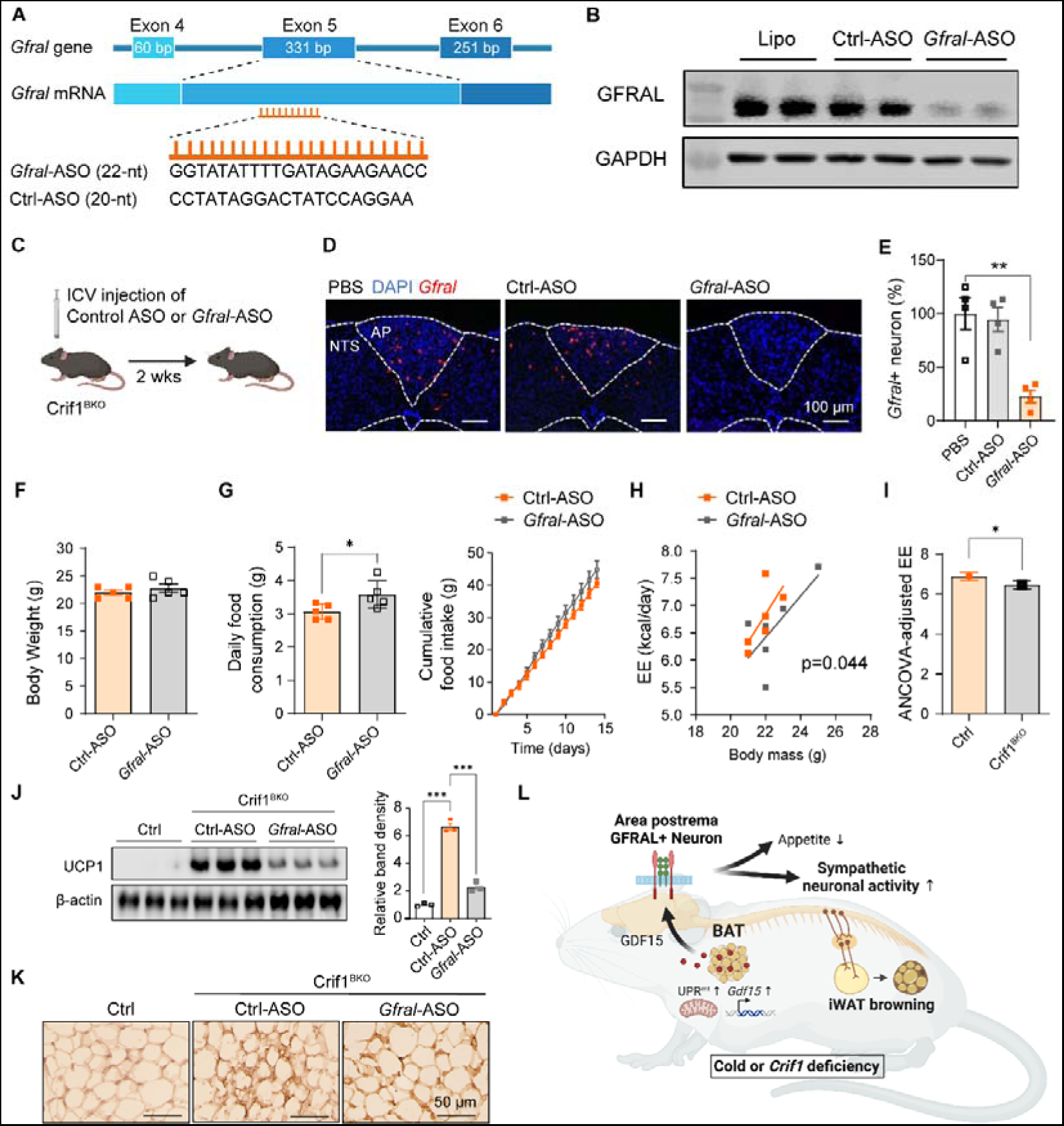
*Gfral* antisense oligonucleotides (ASOs) prevent GDF15-mediated phenotypic changes in Crif1^BKO^ mice. **A**, Design and sequence of ASO targeting *Gfral* mRNA (*Gfral* ASO). The *Gfral* gene in mice include nine exons, and the *Gfral* ASO targets 22 nucleotides within exon 5. **B**, Representative immunoblots for GFRAL 1 week after treatment with lipid-based carriers (lipo), non-targeting control ASOs (Ctrl ASOs), and *Gfral* ASOs. **C**, Intracerebroventricular (ICV) injection of *Gfral* ASO to Crif1^BKO^ mouse. **D**,**E**, *Gfral* mRNA expression 2 week post-injection with PBS, Ctrl ASO, and *Gfral* ASO (**D**), and relative expression of *Gfral*+ neurons in the AP and NTS (**E**). In (**E**), the number of *Gfral*+ neurons were normalized to those in PBS-injected mice. **F–I**, Body weight (**F**) and food intake (**G**) and energy expenditure (EE; **H** and **I**) in Crif1^BKO^ mice, post-injection of Ctrl ASOs and *Gfral* ASOs. **J,K**, UCP1 expression in inguinal white adipose tissue (iWAT) **(J)** and representative images of UCP1-immunostained sections of inguinal WAT **(K)** in control and ASO-pretreated Crif1^BKO^ mice housed at room temperature. Bar graph shows relative protein expression. **L**, Schematic model of the consequences of mitochondrial unfolded protein response (UPR^mt^) activation and *Gdf15* expression in brown adipose tissue (BAT) on systemic energy homeostasis. *, *P* < 0.05; **, *P* < 0.01; ***, *P* < 0.001 by Student’s t-test. Data are represented as mean ± SEM.

We assessed energy expenditure and UCP1 expression in the inguinal WAT of Crif1^BKO^ mice treated with *Gfral* ASOs. These mice exhibited significantly higher food intake than control ASO-treated Crif1^BKO^ mice, although the body weight did not show significant change (**Figs. 7F,G**). Notably, *Gfral* ASO diminished differences in energy expenditure caused by *Crif1* deletion (**Figs. 7H,I**). Furthermore, we observed lower UCP1 expression in the inguinal WAT of *Gfral* ASO-treated Crif1^BKO^ mice compared with control ASO-treated mice (**Fig. 7J**). Histological assessment showed that *Gfral* ASO-injected mice had fewer multilocular adipocytes than control ASO-injected mice (**Fig. 7K**). To identify the role of GFRAL neurons in beige fat formation in mice exposed to cold, we housed GDF15 knockout (KO) mice in a cold environment for 1 days at 7°C and measured UCP1 expression. When we cold-exposed GDF15 KO mice, serum concentrations of GDF15 and GFRAL-expressing neuronal activation in the AP and NTS were significantly lower than cold-exposed WT mice (**Supplementary** Figs. 7I,J). After 24 h of cold exposure, *Ucp1* mRNA expression was significantly lower in the inguinal WAT of GDF15 KO mice than WT mice, while UCP1 protein expression in inguinal WAT tended to be lower in GDF15 KO mice than WT mice (**Supplementary** Figs. 7K–M).

In summary, these results demonstrate that knockdown of *Gfral* in the hindbrain via ASOs prevents the suppression of food intake, along with the elevation in energy expenditure, associated with GDF15 expression. *Gfral* knockdown also results in lower UCP1 expression in inguinal WAT, indicating a diminished thermogenic response. Moreover, the suppression of WAT browning in in cold-exposed GDF15 KO mice confirms the critical role of GFRAL in regulating beige fat formation. These findings underscore the significant impact of hindbrain GFRAL pathways on whole-body energy balance in response to GDF15, which increases with mitochondrial stress in BAT (**Fig. 7L**).

## Discussion

Adaptive thermogenesis in adipose tissue is a key homeostatic mechanism by which animals maintain body temperature in cold environments. Brown adipocytes have developed a range of compensatory mechanisms to offset these deficiencies and sustain thermogenesis; as such, this process persists even when there are defects in the mitochondrial respiratory chain (Masand *et al*., 2018; Pereira *et al*., 2021; Verdeguer *et al*., 2016). For example, in response to mitochondrial dysfunction, brown adipocytes increase UCP1 expression, which elevates metabolic rate and heat production (Masand *et al*., 2018). Additionally, activation of the SNS increases catecholamine release, which stimulates lipid and glucose metabolism for heat production. Alternative pathways, such as ketogenesis, also support thermogenesis when mitochondrial function is compromised (Panic *et al*, 2020). These compensatory mechanisms enable brown adipocytes to maintain thermogenic activity despite mitochondrial dysfunction; however, they may be less efficient and therefore insufficient for maintaining metabolic rate. As a result, communication between brown and white adipose depots is required to offset OXPHOS defects in brown adipocytes. In line with this, previous findings indicate an association between BAT mtDNA-ETC subunit mRNA expression, proteome imbalance, impaired thermogenesis, and reduced adiposity. Likewise, mitochondrial transcription factor A deficiency in BAT is associated with ETC proteome imbalance, defective respiration, and reduced adiposity (Masand *et al*., 2018). However, the precise mechanisms linking mitochondrial stress in BAT to a systemic adaptive metabolic response remains unclear.

Previous studies indicate that GDF15 expression increases in response to cellular stress, such as oxidative stress and mitochondrial respiratory chain dysfunction (Choi *et al*., 2020). Mitochondrial dysfunction typically leads to an increase in oxidative stress, which can trigger the ISR (Mick *et al*, 2020) and lead to subsequent upregulation of GDF15. In addition, ablation of CRIF1 function, specifically in liver and adipose tissue, results in elevated circulating levels of GDF15 (Choi *et al*., 2020; Kang *et al*., 2021) and GFRAL-positive neuron activation in the hindbrain, which coincides with increased UCP1 expression in inguinal WAT. UCP1 transgenic mice that overexpress UCP1 with skeletal muscle specificity also exhibit elevated plasma GDF15 levels and WAT browning (Ost *et al*, 2020). Conversely, GDF15 deficiency leads to diminished inguinal WAT UCP1 expression in these models (Huesing *et al*., 2021; Kang *et al*., 2021). Therefore, GDF15, upregulated by cellular stress, appears to be instrumental in triggering adaptive thermogenesis in WAT. Indeed, several studies show that GDF15 regulates metabolism and increases thermogenesis in WAT when mitochondrial defects in peripheral organs, such as skeletal muscle, exist (Huesing *et al*., 2021; Kang *et al*., 2021).

In line with previous findings, we observed a relationship between cellular stress and metabolic regulation mediated by GDF15. In our study, UPR^mt^ and the ISR were activated in Crif1^BKO^ mice: this was likely responsible for higher BAT GDF15 expression and serum GDF15 concentrations in Crif1^BKO^ mice. The deletion or dysfunction of CRIF1 in tissues typically results in impaired OXPHOS function. This often triggers the UPR^mt^ due to mitochondrial stress from accumulated misfolded proteins (Durieux *et al*, 2011; Nargund *et al*, 2012). While the impairment of BAT OXPHOS is directly linked to CRIF1 deletion, downstream effects, such as protection from obesity and insulin resistance, are consistently observed across different tissues with *Crif1* gene deletion (Choi *et al*., 2020; Chung *et al*., 2017; Kang *et al*., 2021). Furthermore, plasma levels of GDF15 in mice with *Crif1* deletion— whether in skeletal muscle, WAT, or liver—are comparable to those we observed in Crif1^BKO^ mice, levels that stimulate activation of GFRAL neurons in the hindbrain. Therefore, the metabolic benefits of UPR^mt^ activation, combined with the increase in GDF15, appear to result from activation of the GDF15–GFRAL axis in the hindbrain.

The involvement of the GDF15–GFRAL axis in SNS activation has been previously investigated. GDF15-expressing xenografts induce thermogenic gene expression in BAT and lipolytic gene expression in WAT (Chrysovergis *et al*, 2014). GDF15 also stimulates hepatic triglyceride export via β-adrenergic signaling, while chemical sympathectomy completely reverses GDF15-induced body mass loss (Luan *et al*, 2019; Suriben *et al*, 2020). The mechanism behind SNS activation by GDF15 remains elusive, but it may involve hypothalamic neurons in the arcuate nucleus and PVH, which regulate energy homeostasis (Johnen *et al*, 2007). GDF15 might also influence the hypothalamic-pituitary-adrenal axis (Cimino *et al*, 2021): this plays a crucial role in the stress response, although the impact of GDF15 on the hindbrain receptor GFRAL within the context of hypothalamic-pituitary-adrenal axis and proopiomelanocortin neuron activation is yet to be clarified in Crif1^BKO^ mice. Spinal preganglionic neurons, the final controllers of the SNS crucial for environmental stress adaptation, are not believed to express GFRAL (McCarthy, 2006); therefore, activation of these neurons in Crif1^BKO^ mice may be mediated by neuronal or humoral pathways. We confirmed the activation of T7–T10 spinal preganglionic neurons in Crif1^BKO^ mice and identified the responsible neural pathway using retrograde tracing with a pseudorabies virus. This virus traced axons from inguinal WAT to cell bodies in the spinal cord preganglion and hindbrain. Fluorescence microscopy enabled visualization of neurons connected to inguinal WAT. Our results suggest that the GDF15–GFRAL axis forms a unique neural network involving GFRAL-positive neurons, spinal preganglionic neurons, and the end organ, inguinal WAT, which results in beige adipocyte formation in mice with high GDF15 concentrations.

Recent studies have demonstrated the therapeutic potential of activating the GFRAL– SNS through GDF15 administration (Sjøberg *et al*, 2023; Wang *et al*, 2023). Administration of GDF15 not only suppresses appetite but also counteracts reductions in energy expenditure, which leads to more effective weight loss and a greater reduction in non-alcoholic fatty liver disease than caloric restriction alone (Mick *et al*., 2020). The influence of GDF15 on energy expenditure is mediated by a GFRAL–β-adrenergic signaling mechanism that enhances fatty acid oxidation in mouse skeletal muscle. In particular, the GFRAL–β-adrenergic signaling axis increases fatty acid oxidation and stimulates futile calcium cycling in mouse skeletal muscle (Wang *et al*., 2023). Additionally, GDF15 enhances insulin action in obese rodents without the need for weight loss, increasing insulin sensitivity due to decreased glucose production and increased glucose uptake. This insulin-enhancing effect operates through the GFRAL receptor and β-adrenergic signaling, impacting both the liver and adipose tissue (Sjøberg *et al*., 2023). Nonetheless, tissue responses and therapeutic effects of a pharmacological dose of GDF15 vary considerably across species and even among individuals within a species because of genetic, metabolic, and environmental factors (Moon *et al*, 2020). In addition, there are significant differences in GDF15 plasma levels resulting from pharmacological GDF15 administration and those resulting from gene editing, such as in Crif1^BKO^ mice. Although we observed higher GDF15 plasma concentrations in Crif1^BKO^ (about 500-1000 pg/mL) than in control mice, these levels are of a lower magnitude than those resulting from pharmacological GDF15 administration (about 50,000-110,000 pg/mL) (Wang *et al*., 2023), meaning that the energy expenditure seen in the skeletal muscle of mice treated with GDF15 might be less pronounced in Crif1^BKO^ mice. However, even in Crif1^BKO^ mice, we observed lower food intake and body weights and higher energy expenditure than in control mice. These results suggest that even at a subtherapeutic concentration, GDF15 can have a significant impact on appetite and metabolic health.

Cold exposure stimulates mitochondrial biogenesis in brown adipocytes through a complex process involving several signaling pathways. Specifically, it stimulates expression of the transcriptional co-activator PPARGC1A, which regulates mitochondrial biogenesis (Puigserver *et al*, 1998). PPARGC1A activates several transcription factors, including nuclear respiratory factor 1 and mitochondrial transcription factor A. These transcription factors control the expression of genes involved in mitochondrial biogenesis and function in brown adipocyte (Wu *et al*, 1999), which contribute to greater thermogenic capacity of BAT in response to cold exposure. We found that GDF15 expression was induced in the BAT of mice exposed to cold temperatures, and that this was accompanied by increases in the expression of genes involved in mitochondrial biogenesis and the UPR^mt^, as well as mitochondrial chaperones and proteases. The induction of the UPR^mt^ was more marked in differentiated brown adipocytes, which are capable of thermogenesis, than in undifferentiated cells; however, the induction of the UPR^mt^ was less marked under thermoneutral conditions in mice with defective OXPHOS. These results suggest that the induction of the mitochondrial stress response is dependent on the metabolic workload of the cells. This may be analogous to the induction of the ISR by mitochondrial defects dependent on the metabolic state of the cell (Mick *et al*., 2020), and the induction of endoplasmic reticulum stress in WAT resulting from the elevated metabolic workload present in obesity (Ozcan *et al*, 2004; Zhao *et al*, 2002).

In conclusion, GDF15 expression induced by the UPR^mt^ in BAT causes browning of inguinal WAT through the activation of GFRAL-expressing spinal preganglionic sympathetic neurons. These findings outline a compensatory mechanism involving both BAT and WAT that may provide novel targets for therapeutic interventions aimed at increasing energy expenditure in patients with obesity and the metabolic syndrome.

## Materials and Methods

### Animals

To generate brown adipocyte-specific Crif1-deficient mice, Crif1 floxed mice (Kwon *et al*, 2008) were cross-mated with UCP1-Cre mice (Jackson Laboratory, Cat. 024670). C57BL/6J male mice were purchased from Orient Bio (Seongnam, Korea). GDF15 KO mice derived from the inbred C57BL/6 strain were provided by Dr. S. Lee (Johns Hopkins University School of Medicine, Baltimore, MD). All mice had free access to a standard chow diet (ENVIGO, Teklad global 18% protein, Cat. 2918C) and water unless indicated. To induce obesity, mice were fed a high-fat diet (HFD; 60% of total energy intake as fat, ENVIGO, Cat. TD.06414). Animals were housed under controlled temperature conditions (22 ± 1°C) unless indicated and subjected to a 12 h light–dark cycle, with light from 08:00 to 20:00 h. During the cold challenge (5∼7°C), body temperatures of Ctrl or Crif1^BKO^ were monitored with an implantable programmable temperature transponder (Bio Medic Data Systems, Seaford, USA). All animal procedures were approved by the Institutional Animal Care and Use Committee of the Chungnam National University School of Medicine (Daejeon, Korea).

### Metabolic phenotyping

Food intake levels and body weights were monitored weekly after weaning until sacrifice. For the pair-feeding experiment, 6-week-old mice were separately housed and fed an HFD for 18 weeks. Food intake was measured daily, and the average amount of food consumed by the Crif1^BKO^ group was given to the Ctrl-pair-fed group. Body composition (lean mass and fat mass) was measured using dual X-ray absorptiometry (InAnlyzer, MEDIKORS, Seongnam, Korea) during the late light phase at the indicated ages. Energy expenditure were determined using an OxyletProTM system (Panlab, Barcelona, Spain). Mice were placed in the OxyletProTM for 48 h to acclimatize to these conditions before measurement. During this period, day–night cycles were the same as those during the initial housing of the animals and food was provided as pellets on the floor of the OxyletProTM. Data were analyzed using METABOLISM software, version 2.2 (Panlab). For the glucose tolerance test, mice were fasted for 16 h, and then 2 g glucose per kilogram of body weight was injected into the intraperitoneal cavity. The insulin tolerance test was performed by measuring blood glucose after 6 h of fasting followed by intraperitoneal injection of 0.75 U/kg insulin (Humalog Lilly, Indianapolis, IN, USA). Blood samples were obtained from the tail vein for glucose measurement immediately before, and at the indicated times after, injections. Glucose levels were measured using a glucometer (ACCU-CHEK, Roche Diabetes Care, Inc., IN, USA).

### Blood pressure and heart rate measurement

Blood pressure (BP) was measured from 10 to 13-week-old control and Crif1^BKO^ mice weighing 25 g or more using a non-invasive tail cuff system (CODA, Kent Scientific, Torrington, CT). Mice were acclimated to the restraining device by being placed them in an animal holder for 5 minutes for the first 4 days and 15 minutes for the next 3 days. On the day of BP measurement, each animal was allowed to acclimate for 5 minutes in the animal holder at 9 AM. Prior to BP measurement, the temperature of the mice’s tails was verified to be between 32-35°C. BP values were obtained by averaging at least 20 measurements obtained from 20 inflation/deflation cycles. Systolic and diastolic BP and heart rate were determined using the CODA software (Kent Scientific) connected to a computer system.

### Immunostaining of brain and spinal cord sections

Mice were anesthetized with isoflurane and perfused with 20 mL normal saline followed by 20 mL 4% paraformaldehyde via the left ventricle of the heart. Whole brain was collected and fixed with 4% paraformaldehyde for 24 h at 4°C, before being dehydrated in a PBS-based 30% sucrose solution until the tissues sank to the bottom of the container, usually after 48 h at 4°C. Coronal brain sections and transverse spinal cord sections were collected at a 30 μm thickness using a cryostat (Leica, Wetzlar, Germany). Brain slices were blocked with 5% donkey serum in 0.3% PBS-T for 1 h at RT and then incubated with a sheep anti-GFRAL antibody (1:500, Thermofisher, Cat. PA5-47769) and/or a rabbit anti-c-Fos antibody (1:500, Abcam, Cat. ab190289) at 4°C overnight. In the case of spinal cord staining, heat-induced epitope retrieval was required before the blocking step. The heat-induced antigen retrieval method was modified from a reference method (Jiao *et al*, 1999) to use 10 mM sodium citrate buffer and double boil the slides within the buffer using a 60°C water bath. After antigen retrieval, spinal cord slices were blocked in the same way as brain slices and incubated with a goat anti-ChAT antibody (1:100, Chemicon, Cat. AB144P) and a rabbit anti-c-Fos antibody (1:500, Abcam, Cat. ab190289) at 4°C overnight. Slices were then incubated with combinations of secondary antibodies from Invitrogen, including donkey anti-rabbit 647 (1:500, Cat. A31573), donkey anti-goat plus 488 (1:200, Cat. A32814), and donkey anti-sheep 488 (1:500, Cat. A11015), for 1.5 h at RT. For nuclear staining, slides were treated with DAPI (1:10000) for 10 min before mounting. *Z*-stack images were acquired with a confocal microscope (Zeiss, LSM 780), and maximum-intensity *z*-stacks were produced with Zen Black software (Zeiss). Cross-sectional images of the intermediolateral nucleus were taken at 40× magnification, and brain images and regional images of spinal cords were taken at 20× magnification.

### CSF collection for GDF15 quantification

Mice were anesthetized with 2% isoflurane and mounted on stereotaxic frames using ear bars. The mouse snout was tilted down to form an angle of 30–45**°** between the frame base and imaginary line connecting bregma to lambda following the reference method (Šakić, 2019). Next, the scalp and muscle covering the dura mater over the fourth ventricle was gently removed, a 36-gauge beveled needle (WPI, Cat. NF36BV) connected to 1 mL syringe with tubing was slowly inserted into the dura mater, and the CSF was gently aspirated; approximately 8–10 µL of CSF was collected and used for GDF15 quantification.

### Retrograde tracing the neurons innervating inguinal WAT

Mice were anesthetized with 2% isoflurane and the flank was exposed using the lateral decubitus position. PRV-CAG-EGFP (BrainVTA, Cat. P03001) was injected into unilateral inguinal WAT (4 × 200 nL) using a 36-gauge beveled needle (WPI, Cat. NF36BV). After the injection, mice recovered on a heating pad until they were awake and mobile. Mice were sacrificed at 96 and 120 h post-injection, and the brain and spinal cord were isolated to identify sites of EGFP expression and evaluate co-localization of EGFP with ChAT in the spinal cord and GFRAL in the AP/NTS.

### ICV administration of *Gfral* ASOs

Mice at 8–13 weeks of age with average weights of 25–30 g were anesthetized with 3% isoflurane and anesthesia was maintained with 2% isoflurane administered at 0.5 L/min. The head was fixed to a stereotaxic platform (RWD Life Science, China, Cat. 68045). Next, a 36-gauge NanoFil needle was inserted through a burr hole into the right lateral cerebral ventricle using stereotaxic coordinates –0.58 mm posterior from bregma, 1.05 mm lateral from bregma, and 2.2 mm deep from the bregma plane. Injection of a control or *Gfral* ASO (GGTATATTTTGATAGAAGAACC) was performed using a Pump 11 Elite Nanomite pump and Nanomite Injector Unit (Harvard Apparatus; Catalog# 70-4507, Serial# D-301251) at a rate of 2 μL/min. Food intake and body weight measurements, as well as metabolic chamber studies, were performed 1 week postoperatively.

### Western blotting

Mouse tissues were homogenized in lysis buffer (50 mM Tris-HCl, pH 7.4; 150 mM NaCl; 1 mM EDTA, pH 8.0; 0.1% Triton X-100) containing a protease inhibitor cocktail (#11836145001, Roche, Basel, Switzerland) and phosphatase inhibitors (04906837001, Roche) on ice for 30 min using a TissueLyser II (Qiagen, Venlo, Netherlands). After centrifugation at 16,000 × *g* for 15 min, the protein concentrations of the supernatants were measured using a BCA protein assay (#23227, Thermo Fisher Scientific). From each sample, 50 µg protein was loaded onto 8–12% polyacrylamide gels and electrophoresis were performed. The separated proteins were then electrotransferred to 0.45 μm PVDF membranes (#IPVH00010, Millipore) at 200 mA for 2 h. Membranes were blocked with 5% skimmed milk (#T145.2, Roth) in TBS/T buffer (20 mM Tris, 150 mM NaCl, 0.1% Tween 20, pH 7.6) for 1 h and then incubated with primary antibodies overnight at 4°C. After washing three times with TBS/T, the membranes were incubated with secondary antibodies for 1 h at RT and visualized using ECL solution (#34580, Thermo Fisher Scientific). Target protein levels were normalized to those of β-actin, α-tubulin, or glyceraldehyde 3-phosphate. The antibodies used are listed in SupplementaryTable S1.

### Cell cultures

Immortalized brown adipocytes were provided by Dr. Shingo Kajimura (University of California San Francisco, CA). Immortalized brown adipocytes were maintained in Dulbecco’s Modified Eagle’s Medium (DMEM; Welgene, Daegu, South Korea, Cat. LM001-05) supplemented with 10% fetal bovine serum (CYTIVA, Utah, US, Cat. SH30919.03) and 1% penicillin and streptomycin (Welgene, Daegu, South Korea, Cat. LS202-02). Forty-eight hours post-confluence, the cells were differentiated with 0.5 mM 3-isobutyl-1-methylxanthine, 1 μM dexamethasone, 1 µg/mL insulin, 1 nM T3, and 1 µM rosiglitazone. Two days later, the medium was replaced with DMEM containing 10% FBS, 1 nM T3 and 1 µg/mL insulin, and was changed every 2 days for a total of 7 days to achieve complete differentiation. The cells were treated with CL 316,243 (1 µM) for 1.5, 3, or 6 h, or oligomycin (Sigma Aldrich, Cat. O4876, 10 µg/mL), CCCP (Sigma Aldrich, Cat. C2759, 2 µg/mL), rotenone (Sigma Aldrich, Cat. R8875, 1 µM/mL), or gamitrinib (Legochem Biosciences, 1 or 2 µM) for 24 h at 37°C.

### RNA sequencing (RNA-seq)

RNA was isolated from the BAT of 8-week-old control and Crif1^BKO^ mice. To construct cDNA libraries, 1 μg RNA and a TruSeq RNA Library Prep Kit v2 (RS-122-2001, Illumina, San Diego, CA, USA) were used, and the results were quantified by qPCR using an 2100 Bioanalyzer (Agilent Technology Inc., Santa Clara, CA, USA). The libraries were used for 100 nt paired-end sequencing by an Illumina HiSeq4000 (Illumina). After removing the low-quality and adapter sequences using Trimmomatic, the reads were aligned with the Mus musculus genome (mm10) using HISAT (ver. 2.0.5). Two types of indexes were used for alignment (a global, whole-genome index and tens of thousands of small local indexes), which were downloaded from the UCSC table browser (http://genome.uscs.edu). StringTie (ver. 1.3.3b) was used to assemble the transcript, and provided the relative abundance estimates as fragments per kilobase of exon per million fragments mapped (FPKM) values of the transcript or gene. After excluding the genes with one more than zero FPKM values, the signal value (FPKM+1) was transformed to a base 2 logarithm and normalized by quantile normalization methods to reduce systematic bias. These values were used for the analysis of differentially expressed genes in the mouse groups. The statistical significance of differentially expressed gene values was calculated using independent t-tests (*P*<0.05) and fold change (|FC|≥2). The false discovery rate, which estimates the frequency of type I statistical errors, was determined by adjusting the *P*-value using the Benjamini and Hochberg algorithm.

### mRNA expression measurement

Tissues were homogenized using a TissueLyser II (Qiagen) and RNA was isolated using TRIzolTM Reagent (Life Technologies, Thermo Fisher Scientific, Cat. 15596018). Complementary DNA(cDNA) was synthesized from 5μg of total RNA using Oligo(dT)_15_ Primer (C1101, Promega, Madison, WI, USA) and M-MLV Reverse Transcriptase (Thermo Fisher Scientific, Cat. 28025). The produced cDNA was then amplified on a 7500 Fast Real-Time PCR System (Applied Biosystems, Carlsbad, CA). Real-time PCR was performed using SFC Green Fast qPCR master mix w/ Low ROX(2x) (SFC, South Korea, Cat.ULH1I301). All quantitative calculations were performed using the ^ΔΔ^CT method. All mouse primer sequences are listed in Supplementary Table S2.

### Enzyme-linked immunosorbent assay

Cell supernatant was obtained from primary hepatocytes by centrifugation at 16,000 × *g* for 5 min. Blood was collected from a retro-orbital sinus, incubated at RT for 2 h and then centrifuged at 600 × *g* for 5 min to obtain serum or CSF. GDF15 (R&D Systems, Minneapolis, MN, USA, Cat. MGD150) and FGF21 (R&D Systems, Cat. MF2100) concentrations were measured using ELISAs according to the manufacturer’s instructions.

### Statistical analysis

Statistical analyses were performed using SPSS software, version 26.0 (IBM, Armonk, NY, USA). Data are reported as mean ± standard errors of the mean (SEMs). A two-tailed Student’s t-test was used to determine differences between the groups. A *P*-value <0.05 was considered statistically significant.

## Data Availability

RNA-Seq data associated with this study are available from Gene Expression Omnibus [GSE233960] (https://www.ncbi.nlm.nih.gov/geo/query/acc.cgi?acc=GSE233960).

## Author Contributions

Conceptualization, J.H.L., J.W.S., and M.S.; Methodology, J.H.L., J.E.K., S.H.J., M.H.L., H.J.H., Z.B., Y.E.K., J.T.K., H.S.Y., and J.K.; Formal Analysis, J.H.L., J.E.K., S.H.J., M.H.L., S.J., and S.E.L.; Investigation, J.E.K., S.H.J., M.H.L., H.J.H., U.Y., Y.S.Y. and M.P.; Writing – Original Draft, J.H.L., J.E.K., and S.H.J.; Writing – Review & Editing, J.H.L., J.A., J.W.S., and M.S.; Supervision, M.S.; Funding Acquisition, J.H.L. and M.S. All the authors read and edited the manuscript, and approved the final version.

## Acknowledgments

This study was supported by a National Research Foundation of Korea (NRF) grant funded by the Korea government, Ministry of Science and ICT (RS-2023-00210819) and NRF-2023R1AC3003438, South Korea, the European Research Council (ERC-AdG-787702), and Global Research Laboratory Program through the NRF of Korea (NRF 2017K1A1A2013124).

