## Supplementary figures for "Adaptive thermogenesis is mediated by GDF15 via the GFRAL neuronal axis in mice"

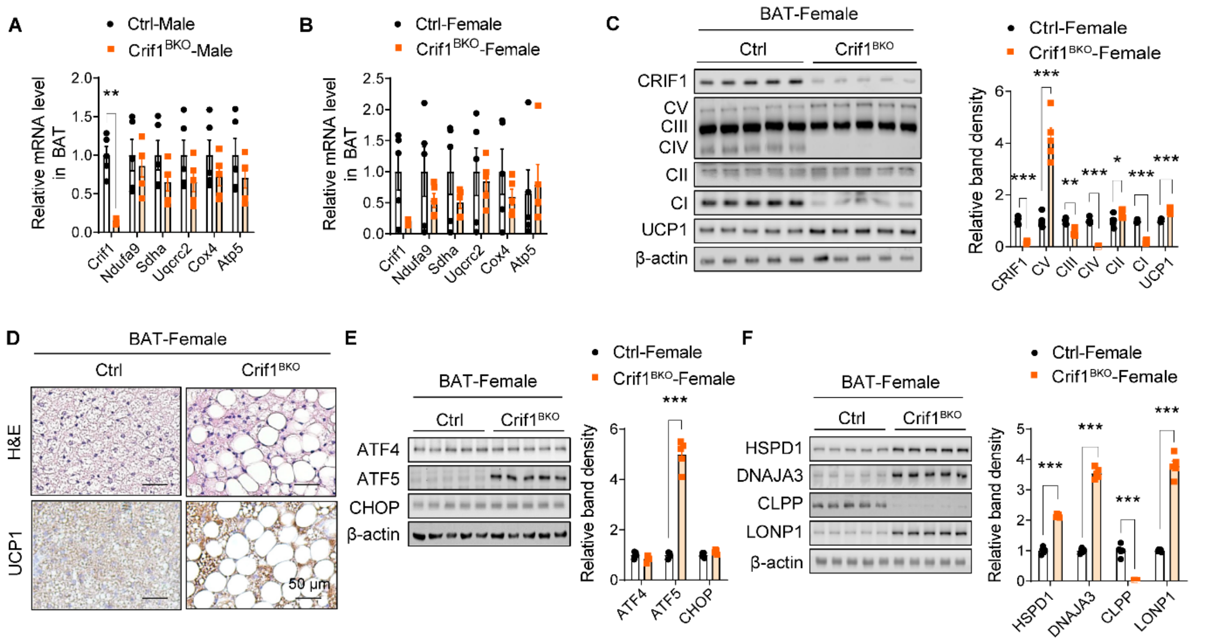


**Figure S1. Cirf1$^{BKO}$ mice show OXPHOS detect and increased expression of UPR$^{mt}$ proteins in brown adipocyte. Related to Figure 1**

**(A, B)** Relative mRNA levels of *Crif1* and OXPHOS in BAT of Ctrl and Crif1$^{BKO}$ mice, (A) for male and (B) for female mice (n=5 per group). **(C)** Immunoblots and band densities for CRIF1, OXPHOS proteins, and UCP1 in the BAT of female Ctrl and Crif1$^{BKO}$ mice. **(D)** Hematoxylin and eosin (H&E)-stained and UCP1 immunostained sections of BAT of female mice. Bar, 50 μm. **(E-F)** Immunoblots and band densities for transcription factors (E) and mitochondrial chaperones and proteins (F) in female mice. *, $P < 0.05$; **, $P < 0.01$; ***, $P < 0.001$ *vs*. Ctrl by Student's t test. Data are represented as mean ± standard error of mean (SEM).

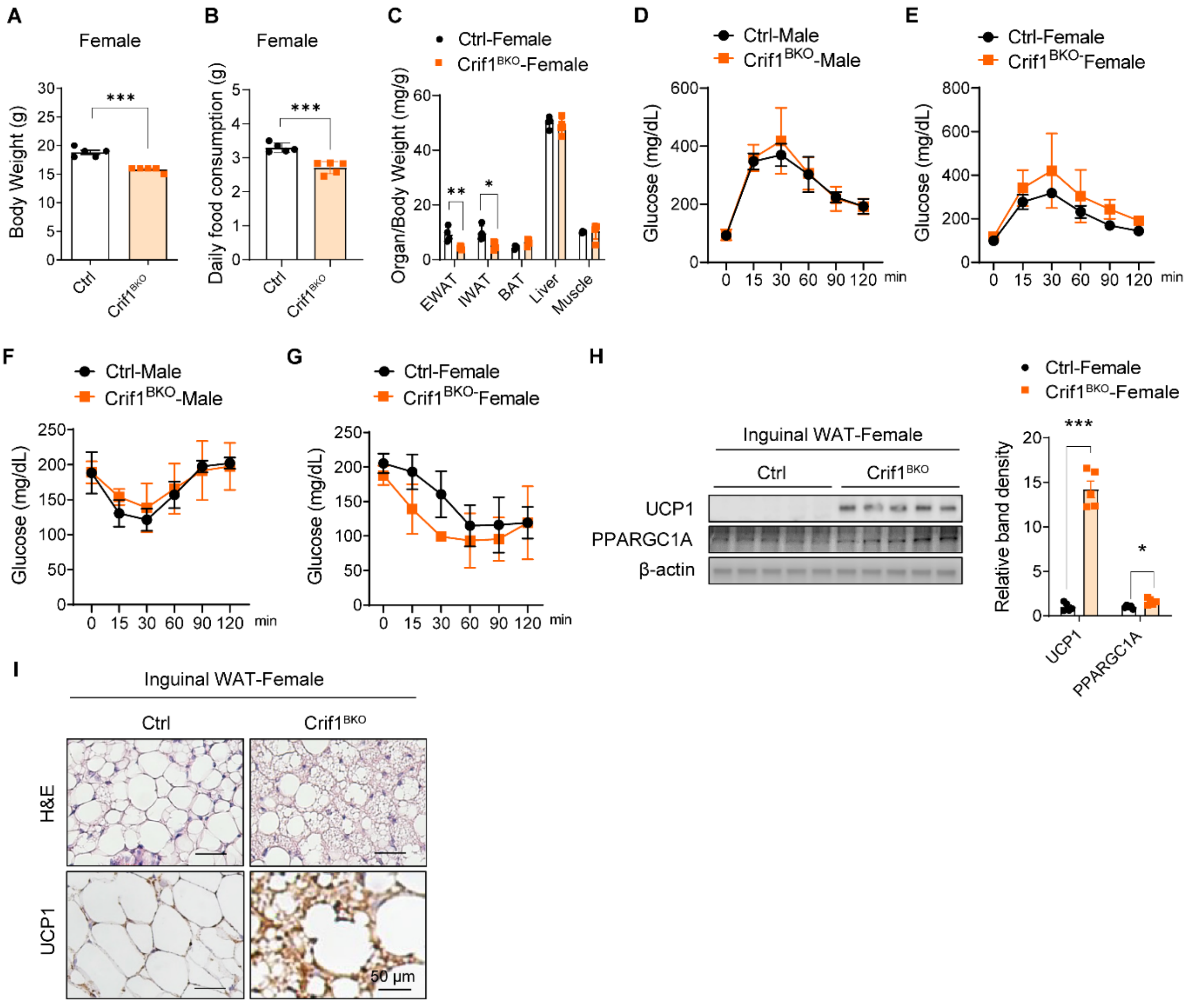


**Figure S2. Metabolic phenotype analysis of mice with Crif1 knockout in brown adipocytes. Related to Figure 2**

**(A-C)** Body mass (A), daily food consumption (B), and organ/body mass ratio (C) for female Ctrl and Crif1$^{BKO}$ mice (n=5 per group). (D) Body temperatures of male Ctrl and Crif1$^{BKO}$ mice housed at room temperature (22–24°C) following cold exposure at 5°C for 1 week (n=4 per group). **(E-F)** Intraperitoneal glucose tolerance test in 8-9 weeks old male (E) and female (F) Ctrl and Crif1$^{BKO}$ mice fed normal chow diet (n=3-4 per group). **(G-H)** Intraperitoneal insulin tolerance test in 9-10 weeks old male (G) and female (H) Ctrl and Crif1$^{BKO}$ mice fed normal chow diet (n=3-4 per group). **(I)** Immunoblots and band densities for UCP1 and PPARGC1A in inguinal WAT of female mice. **(J)** Representative images of H&E-stained and UCP1-immunostained sections of inguinal WAT of female mice. Bar, 50 μm. *, $P < 0.05$; **, $P < 0.01$; ***, $P < 0.001$ *vs*. Ctrl by Student's t test and two-way ANOVA for Figs. S2E-H. Data are represented as mean ± SEM.

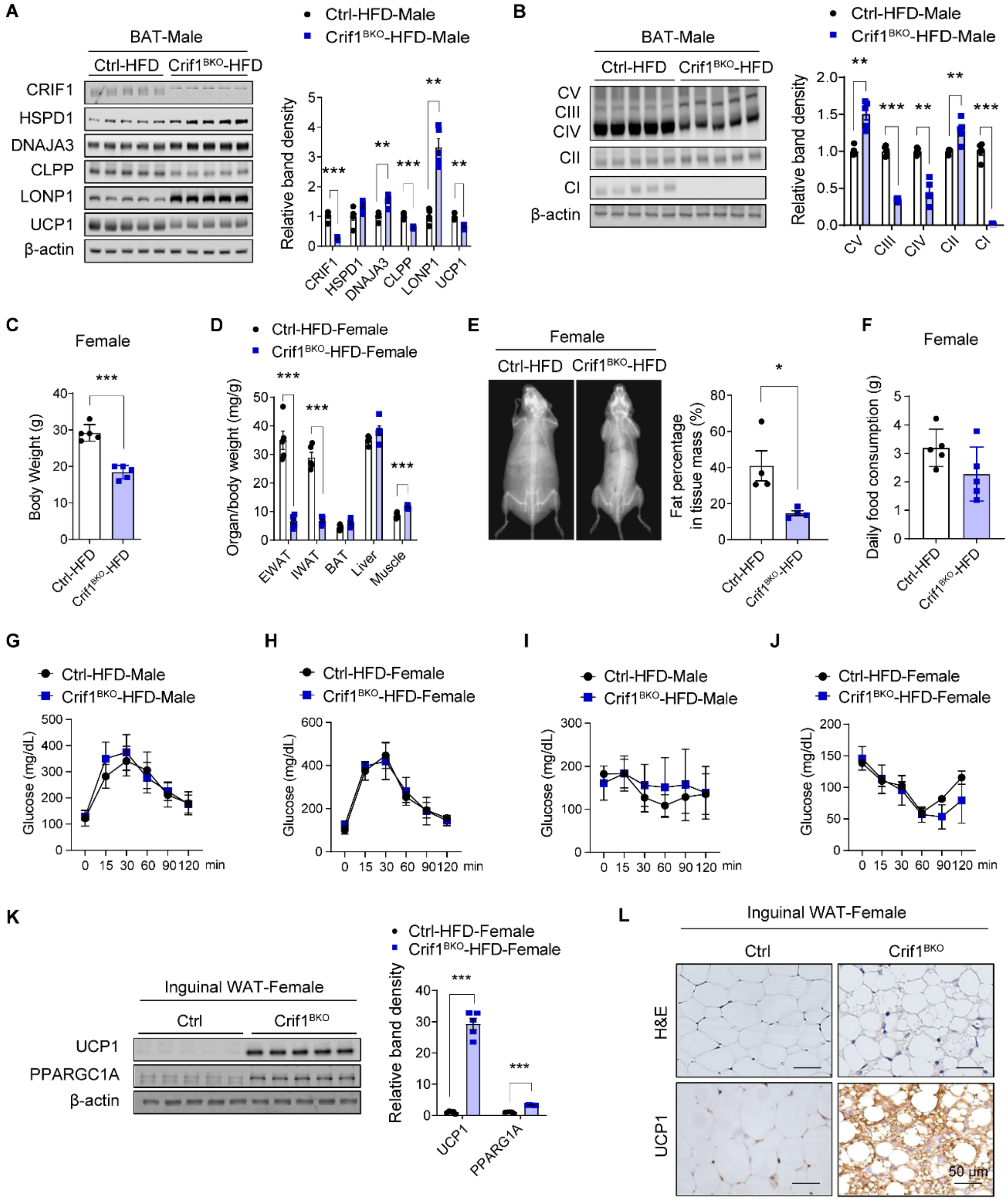


**Figure S3. OXPHOS level of BAT and metabolic phenotype of BAT-specific Crif1 knockout mice with high-fat diet. Related to Figure 2**

**(A)** Immunoblots and its band density of OXPHOS in BAT. **(B-D)** Body masses (B), organ/body mass ratios (C), and fat percentages (D), according to densitometry of female mice. **(E)** Daily food intake of female Ctrl and Crif1$^{BKO}$ mice fed a HFD for 8 weeks (n=5 per group). **(F-G)** Results of intraperitoneal glucose tolerance testing of 14–15-week-old male (F) and female (G) Ctrl-HFD and Crif1$^{BKO}$-HFD mice fed an HFD for 10 weeks (n=5 per group). **(H-**

**I)** Results of intraperitoneal insulin tolerance testing of 13–14-week-old male (H) and female (I) Ctrl-HFD and Crif1$^{BKO}$-HFD mice fed a HFD for 9 weeks (n=5 per group). **(R)** Immunoblots and band densities for UCP1 and PPARGC1A in the inguinal WAT of female mice. **(T)** Representative images of H&E-stained and UCP1-immunostained sections of inguinal WAT of female mice. Bar, 50 µm. *, $P < 0.05$; **, $P < 0.01$; ***, $P < 0.001$ *vs*. Ctrl-HFD by Student's t test and two-way ANOVA for Figs. S3F-I. Data are represented as mean ± SEM.

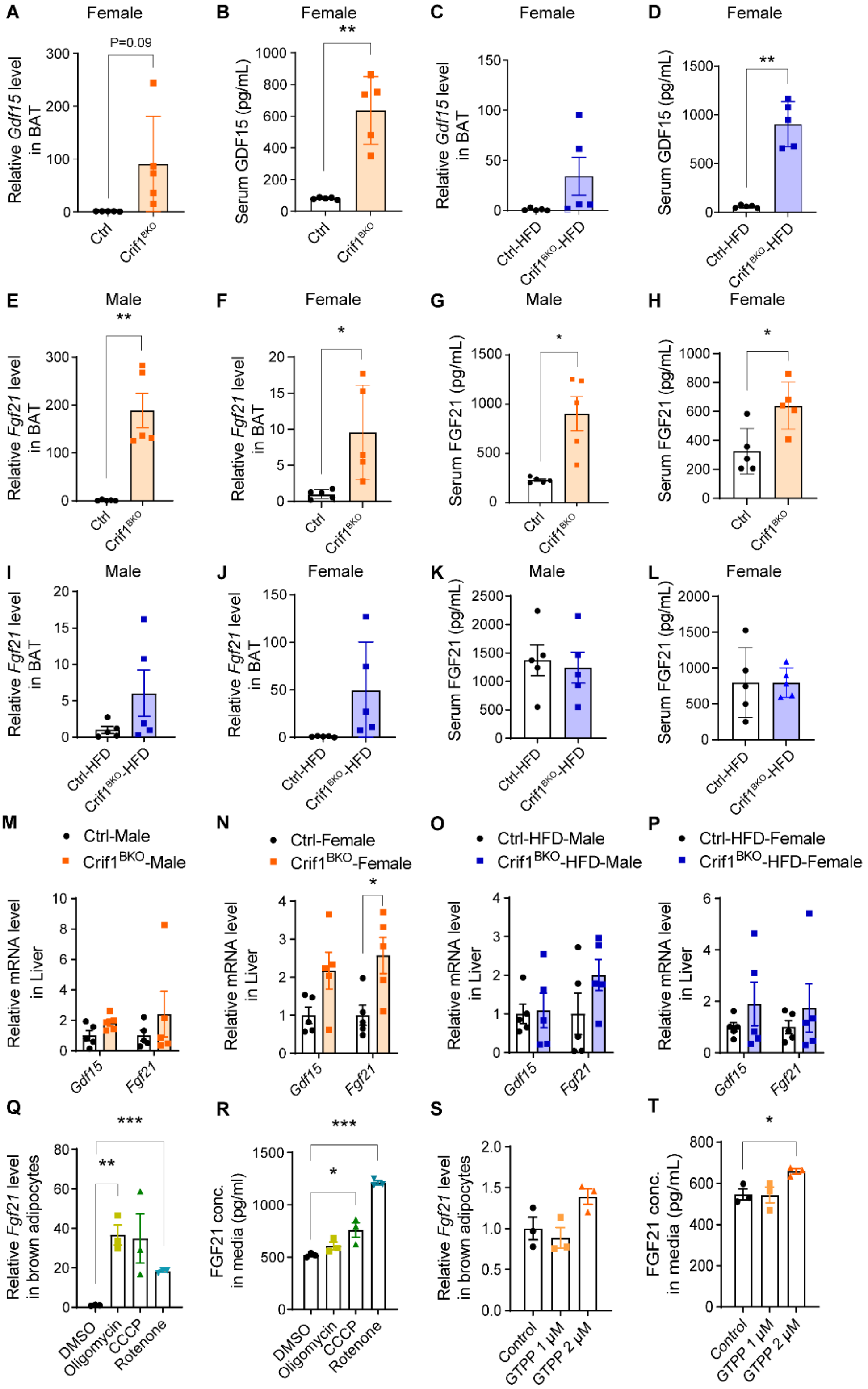


**Figure S4. Mitokine expression in BAT and liver by Crif1 deletion. Related to Figure 2–3**
**(A-B)** Relative *Gdf15* expression in the BAT (A) and serum GDF15 concentration (B) of female Ctrl and Crif1$^{BKO}$ mice (n=5 per group) characterized in **Figures 2A-H**. **(C-D)** Relative

*Gdf15* expression in the BAT (C) and serum GDF15 concentration (D) of female Ctrl-HFD and Crif1$^{BKO}$-HFD mice characterized in **Figures 2I-O**. **(E-F)** Relative *Fgf21* level in BAT of Ctrl and Crif1$^{BKO}$ mice from **Figure 2**. **(G-H)** Serum FGF21 concentration in Ctrl and Crif1$^{BKO}$ mice. **(I-J)** Relative *Fgf21* level in BAT of Ctrl-HFD and Crif1$^{BKO}$-HFD mice from Figure 2. **(K-L)** Serum FGF21 concentration in Ctrl-HFD and Crif1$^{BKO}$-HFD mice. **(M-N)** Relative *Gdf15* and *Fgf21* levels in the liver of male (M) and female (N) Ctrl and Crif1$^{BKO}$ mice with normal chow diet from **Figures 2A-H and Figures S2.** (**O-P)** Relative *Gdf15* and *Fgf21* levels in the liver of male (O) and female (P) Ctrl and Crif1$^{BKO}$ mice fed with HFD from **Figures 2I-O and Figures S3. (Q)** Relative *Fgf21* levels in differentiated brown adipocytes treated with OXPHOS inhibitors; Oligomycin 10ug/ml, CCCP 2ug/ml, Rotenone 1uM/ml for 24h. **(R)** FGF21 conc. in media from **Figure S4Q**. **(S)** Relative *Fgf21* levels in differentiated brown adipocytes treated with gamitrinib for 24h. **(T)** FGF21 conc. in media from **Figure S4S**. *, P < 0.05; **, P < 0.01; ***, P < 0.001 *vs*. Ctrl, Ctrl-HFD, or DMSO.

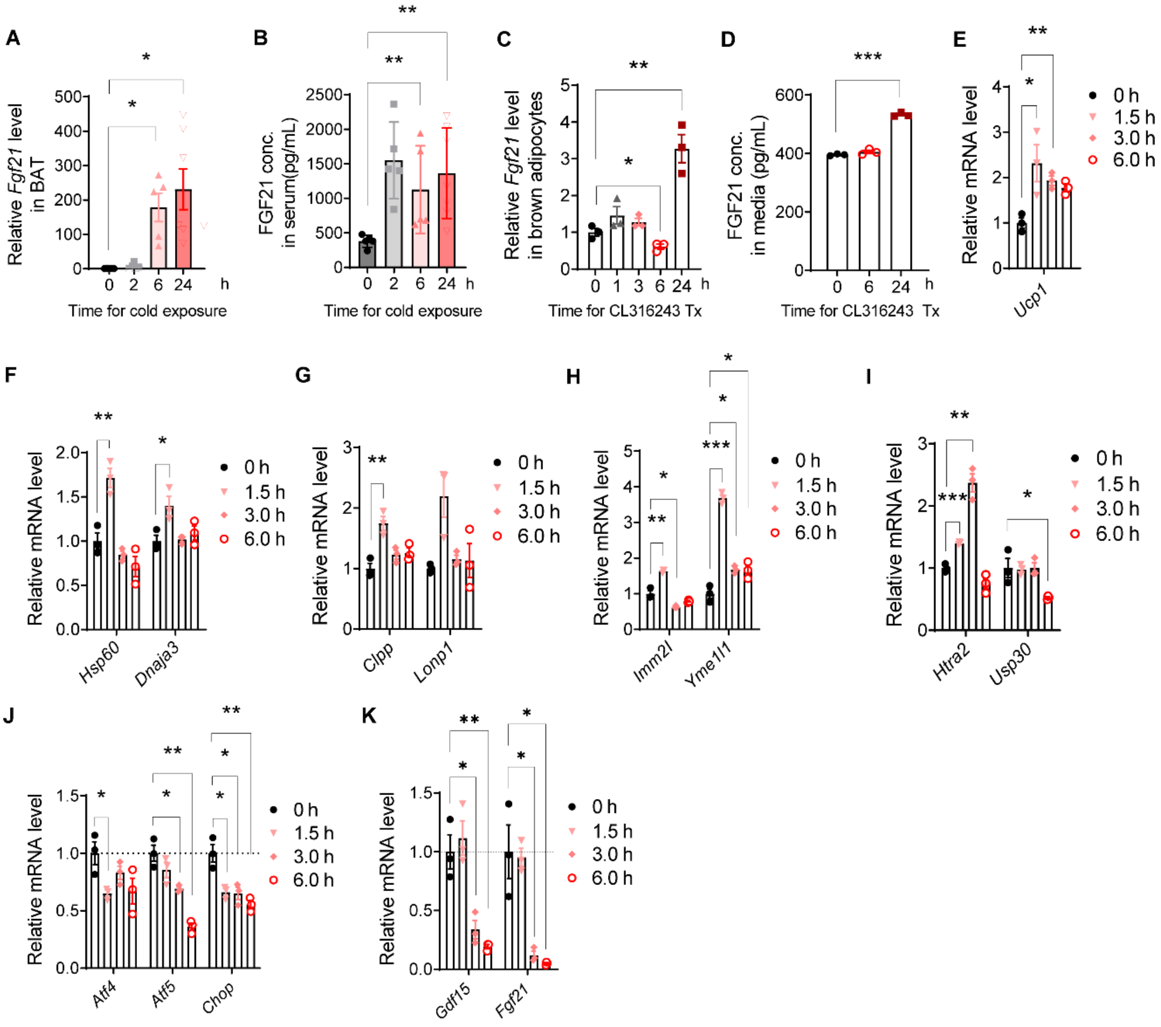


**Figure S5. Mitokine expression by cold-exposure. Related to Figure 4**

**(A-B)** Relative *Fgf21* level of BAT (A) and serum FGF21 concentration (B) of cold-exposed mice **(C-D)** Relative *Fgf21* level (C) and FGF21 concentration in media (D) from differentiated immortalized brown adipocytes treated with 1μM CL316,243. **(E-I)** Relative mRNA levels of *Ucp1* (E) and mitochondrial chaperones (F) and proteases (G-I) in undifferentiated immortalized brown adipocytes treated with 1μM CL316,243. **(J-K)** Relative mRNA level of transcription factors (J), *Gdf15* and *Fgf21* level (K) in undifferentiated immortalized brown adipocytes treated with 1μM CL 316,243. *, P < 0.05; **, P < 0.01; ***, P < 0.001 *vs*. 0h.

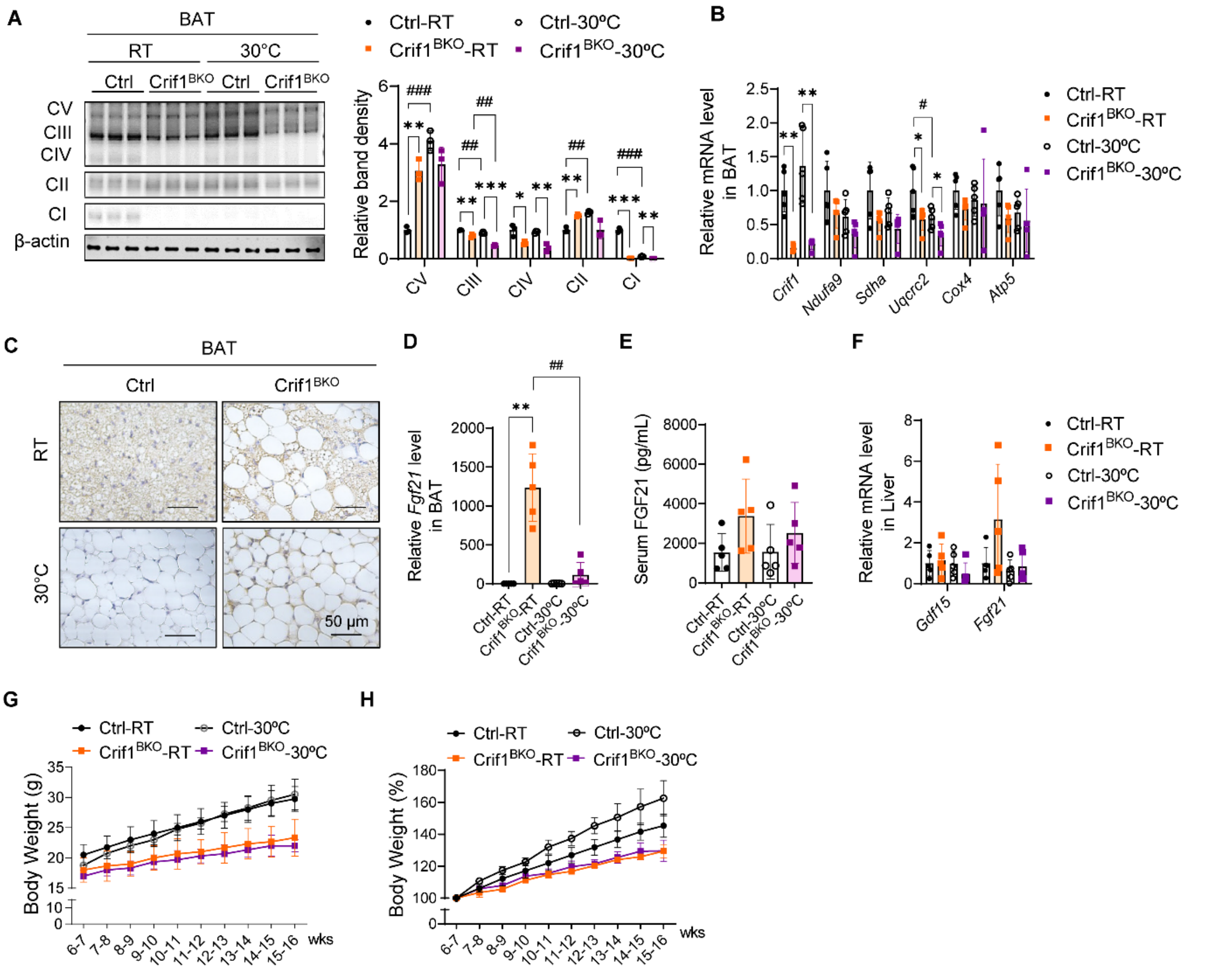


**Figure S6. OXPHOS and mitokine expression regarding to the thermoneutral condition in the BAT-specific Crif1 knockout mice. Related to Figure 5**

**(A)** Immunoblots and its band density of OXPHOS in BAT of 12-13 weeks old male Ctrl and Crif1$^{BKO}$ mice housing room temperature (22~24°C) (Ctrl-RT or Crif1$^{BKO}$-RT) or 30°C (Ctrl-30°C or Crif1$^{BKO}$-30°C) for 8wks **(B)** Relative mRNA level of OXPHOS genes in BAT. **(C)** Representative images of anti-UCP1 staining of BAT. bar, 50 ㎛. **(D-E)** Relative *Fgf21* level in BAT (D) and serum FGF21 concentration (E) in Ctrl and Crif1$^{BKO}$ mice housing in RT or 30°C. **(F)** Relative mRNA level of *Gdf15* and *Fgf21* level in liver. **(G-H)** Body weight (G) and its percentage compared to 6-7 wks, the start of housing condition (H) in 15-16 weeks old male Ctrl and Crif1$^{BKO}$ mice housing room temperature (22~24°C) (Ctrl-RT or Crif1$^{BKO}$-RT) or 30°C (Ctrl-30°C or Crif1$^{BKO}$-30°C) for 9wks (n=3-4 per group) *, $P < 0.05$; **, $P < 0.01$; ***, $P < 0.001$ *vs*. Ctrl. #, $P < 0.05$; ##, $P < 0.01$; ###, $P < 0.001$ *vs*. RT.

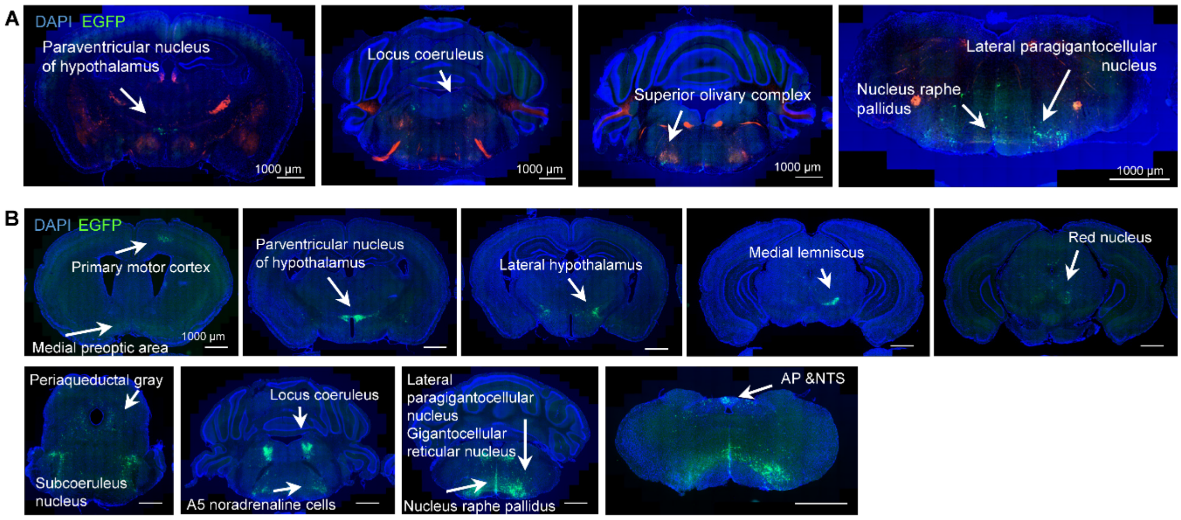


C

| WT1 120hr | total GFRAL+ | GFRAL+ and EGFP+ | GFRAL+ EGFP+ / GFRAL+ |
|---|---|---|---|
| slice 1 | 54 | 11 | 20.4 |
| slice 2 | 53 | 3 | 5.7 |
| slice 3 | 34 | 3 | 8.8 |
| slice 4 | 12 | 1 | 8.3 |
| Sum | 153 | 18 | 11.8 |

| WT2 120hr | total GFRAL+ | GFRAL+ and EGFP+ | GFRAL+ EGFP+ / GFRAL+ |
|---|---|---|---|
| slice 1 | 24 | 10 | 41.7 |
| slice 2 | 49 | 15 | 30.6 |
| slice 3 | 48 | 10 | 20.8 |
| slice 4 | 12 | 2 | 16.7 |
| Sum | 133 | 37 | 27.8 |

Slice 1 Rostral
↑
↓
Slice 4 Caudal

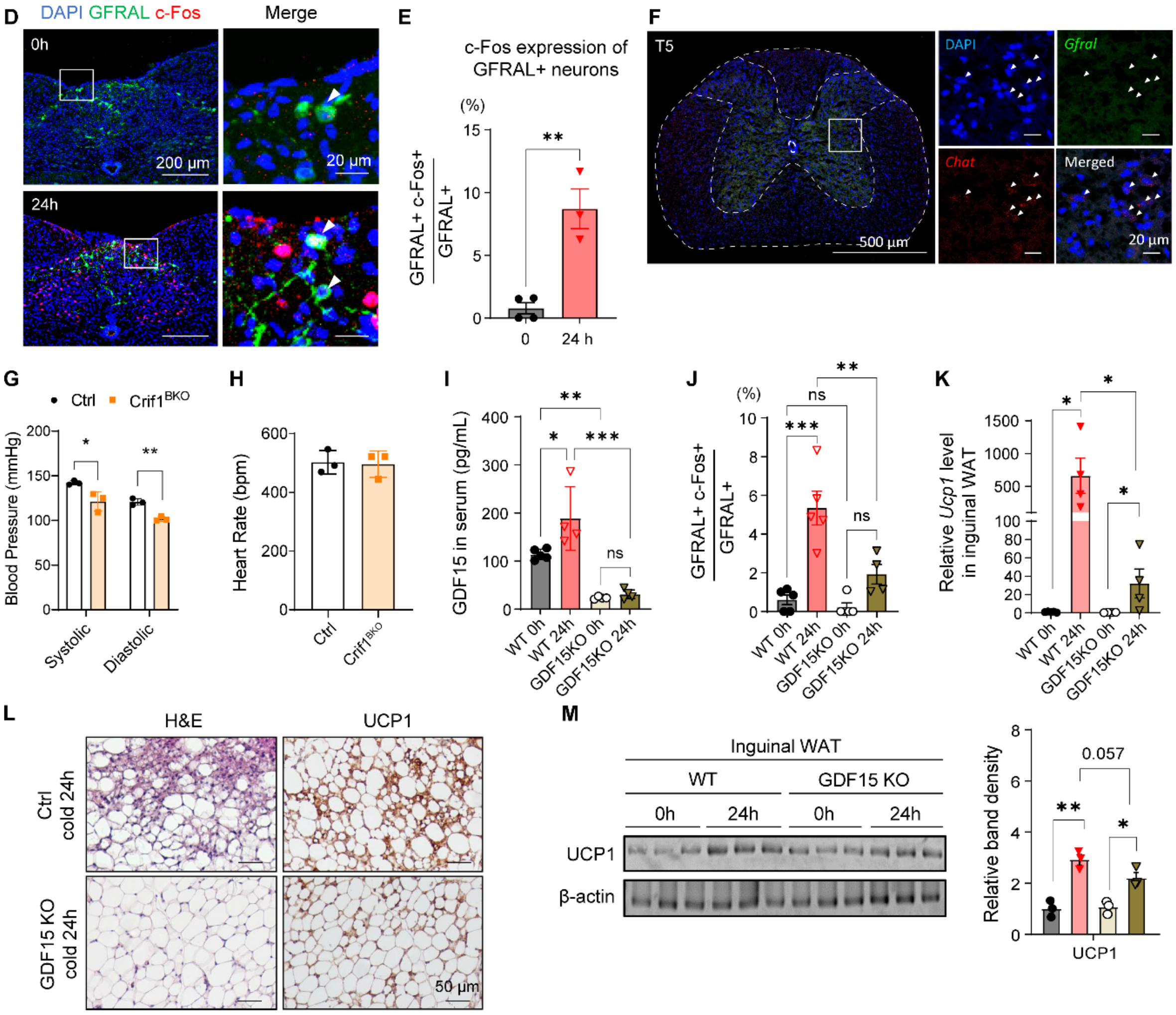

**Figure S7. Cold responsive GFRAL-expressing neurons and their connection with inguinal WAT. Related to Figure 6**

**(A)** Representative images of immunofluorescence (IF) of the brain of ChAT-IRES-Cre::tdTomato mice, after 96h of EGFP-expressing pseudorabies virus injection into the unilateral inguinal WAT. **(B)** Representative images of IF of the brain of 10 weeks old wild-type C57BL/6 male mice after 120h of EGFP-expressing pseudorabies virus injection in the unilateral inguinal WAT. **(C)** EGFP expression among GFRAL-expressing neurons in AP and NTS of four slices from rostral to caudal direction from **Fig. 6J**. **(D-E)** Representative images of FOS expression of GFRAL-expressing neurons in AP and NTS before and after 24h of cold exposure (D) and quantification of FOS expression (E). **(F)** RNAscope *in situ* hybridization depicting *Gfral* and *Chat* in T5 spinal cord section. **(G-H)** Blood pressure (G) and heart rate (H) measured in tail of 10-11 weeks old male Ctrl and Crif1$^{BKO}$ mice (n=3 per group). (**I-J**) Serum GDF15 concentration (I) and FOS expression of GFRAL-expressing neurons in AP and NTS (J) before and after 24h of cold exposure in wild-type and GDF15 KO mice. **(K-M)** Relative *UCP1* level in inguinal WAT (K), representative images of H&E-stained and UCP1-immunostained sections of inguinal WAT (L), and immunoblots and its band density of UCP1 in inguinal WAT (M) before and after 24h of cold exposure in Ctrl and GDF15 KO mice. *, P < 0.05; **, P < 0.01; ***, P < 0.001; ****, P < 0.0001 *vs*. 0h.
